# A spatial and single-cell transcriptomic atlas of the embryonic synovial joint and its explant model

**DOI:** 10.64898/2026.10.19.760178

**Authors:** Sharon E.S. Onggo, Ji-Hye Yea, Kaily K. Young, Kaiwen Chen, Haneul Cho, Zhirui Ning, Patrick Cahan

## Abstract

Synovial joint formation requires the coordinated differentiation of multiple cell types while balancing opposing migratory cues that recruit joint progenitors and exclude other populations. More specifically, a subset of limb bud mesenchyme is recruited into the interzone while neighboring chondrocytes and endothelial cells are excluded. Existing single-cell references sample this process sparsely, without position, or without the surrounding mesenchyme. Here, we present an atlas of the mouse hindlimb from E11.5 to E15.5 that pairs 228,703 single-cell transcriptomes with 47 spatial transcriptomic sections, annotates 56 cell types by their location, and extends the same profiling to limb explant culture. Explants preserve the lineages and axial patterning of the limb for three days, with delayed chondrocyte hypertrophy. Using the atlas, we found that Sema3c is co-expressed with Sox9 and Gdf5 during condensation and is required for chondrogenesis and joint formation in culture; that interzone cells carry a chondrogenic signature early and a tenogenic signature late; and that a core interzone signature includes eleven previously unreported markers, one of which, Igfbp5, is required for knee formation in explants.

## INTRODUCTION

Synovial joints enable near-frictionless movement between adjacent bones and are composed of several distinct tissues: a fluid-filled cavity lined by a secretory membrane and enclosed by a fibrous capsule, articular cartilage, and accessory structures such as menisci, tendons, and ligaments. Generating this architecture from a seemingly continuous precartilaginous template entails the coordinated differentiation of multiple lineages. Directed cell migration also plays a crucial role, both by recruiting progenitors to the nascent joint and by excluding some neighboring populations from it.

Initially, the limb forms as a bud of loosely packed, mesenchymal cells interspersed with endothelial cells that form a primitive vascular plexus and covered by an ectodermal epithelial jacket^1,2^. Centrally located mesenchymal cells upregulate Sox9, condense, exclude vasculature^1–3^, and initiate chondrogenesis to form the cartilage templates that presage future skeletal elements^4–6^. Most chondroprogenitors subsequently undergo endochondral ossification to form bone. However, the limb bud mesenchyme (LBM), which is derived principally from lateral plate mesoderm (LPM), also gives rise to the perichondrium and periosteum, tendons and ligaments, joint capsule and synovium, muscle connective tissue, and dermis^7,8^. Several other lineages are immigrants. Myogenic progenitors delaminate from the somites and migrate into the bud, where they are organized into dorsal and ventral muscle masses by the LPM-derived connective tissue around them^9,10^. Somite-derived angioblasts contribute to the vascular plexus^11^; Schwann cells of the peripheral nervous system and melanocytes of the skin derive from migratory neural crest cells^12^. A central aim of the limb development field has been to learn how the multipotent LBM coordinates its own differentiation with the patterning of differentiation of these distinct immigrating populations to form a functional limb.

Within the LBM, fate is determined largely by position. Sox9⁺ cells at the core of each condensation become chondrocytes, whereas cells at its margins adopt perichondrial and, further out, tendon (Scx⁺) and muscle-connective-tissue (Osr1⁺/Tcf4⁺) fates^13–15^; at the cartilage–tendon interface, Sox9⁺/Scx⁺ bipotent progenitors give rise to the enthesis^13,14^. Joint formation is heralded by the interzone (IZ), a morphologically distinct band of flattened, closely apposed mesenchymal cells that appears either within a continuous cartilage template (e.g., knee, elbow, wrist, ankle) or between adjacent cartilage anlagen (e.g., interphalangeal joints)^16^. IZ cells are marked by expression of the TGF-beta-family ligand Gdf5^17,18^. Classic microdissection experiments in chick showed that excising a 100 to 150 um slice containing the IZ causes joint fusion^19–21^, establishing the IZ and its immediately flanking mesenchyme as the source of synovial joint lineages and suggesting that the IZ serves as a transient reservoir of joint progenitors. Consistent with this, genetic lineage tracing has shown that the IZ is populated not in a single event but by continuous recruitment of Sox9⁺/Gdf5⁻ cells from the flanking mesenchyme over roughly three days, while chondrocytes of the adjacent skeletal elements are excluded^22–25^. Time of arrival predicts fate: early Gdf5-expressing cells preferentially populate the epiphysis, whereas later arrivals contribute to the articular cartilage and fibrous components of the joint^22–25^. These observations frame several open questions. First, which cells in the flanking mesenchyme are competent to enter the IZ, and how do they differ from neighbors that remain chondrogenic? Second, when, relative to condensation, does a joint-forming identity first become detectable, and what distinguishes it from the Gdf5⁺ state that follows? Third, beyond Gdf5, which genes define the IZ and demarcate its boundaries? Finally, can any in vitro system reproduce this sequence well enough to interrogate it experimentally? Answering these questions requires a census of LBM states that spans the full window from condensation to cavitation and that is anchored to position within the tissue.

Single-cell transcriptomics is well suited to a census of this kind, and it has been applied to the developing limb in several complementary ways. Unbiased atlases have surveyed the chick autopod^26^, the mouse hindlimb from E11.5 to E18.5^27^ and forelimb from E10.5 to E14.5^28^, and the human limb from post-conception weeks 5–9^29^; ten public mouse datasets spanning limb induction to adult bone have since been integrated into a 133,000-cell meta-atlas of the skeletal lineage^30^. These atlases consistently recover the major lineages — chondrocytes, osteoblasts, tenocytes, fibroblasts, myogenic cells, endothelium, immune cells, and epithelium — and have proposed subdivisions of the mesenchyme that are not evident histologically, including naïve limb progenitors at the limb margin that give rise to separate proximal and autopodial progenitor pools^28^ and, in human, perineural fibroblasts and multiple distinct mesenchymal states^29^. Lineage-enriched studies have resolved heterogeneity that is diluted in whole-limb samples. Profiling of Gdf5-lineage cells from the knee between E12.5 and E15.5 defined the transcriptional paths from IZ to articular cartilage, meniscus, ligament, and synovium, and identified lineage-biased subpopulations as early as E12.5, before a morphologically distinct IZ is apparent^31^. Tracing of Hic1⁺ cells revealed a rare, somite-derived mesenchymal progenitor that enters the limb at E11.5 and contributes not to the primary cartilage anlagen but to fibroblasts, entheses, myotendinous junctions, and other bone superstructures; these cells do not contribute to the primary cartilage anlagen^32,33^. Perturbation studies have used scRNA-seq to link cell-state changes to joint phenotypes. IZ cells are distinguished from flanking chondrocytes by elevated expression of glycolytic genes, and deletion of the glucose transporters Glut1 and Glut3 in preskeletal mesenchyme or in chondrocytes blocks IZ formation in a dose-dependent manner^34^. Conversely, a PDGFR-alpha gain-of-function allele permits initial Gdf5 expression at E12.5 but not its maintenance, redirecting IZ progenitors toward a fibrocartilage fate and fusing the knee^35^. Finally, spatially resolved approaches have begun to add positional information to these transcriptional states. Indexing dissociated cells to their site of origin in E10.5 to E11.5 forelimb buds showed that the positional and cell-type components of identity are encoded by separable gene programs^36^. Spot-based spatial transcriptomics placed scRNA-seq-defined mesenchymal states in tissue context in the human limb^29^; and whole-embryo spatial datasets include limb sections at several stages^37,38^.

Despite this growing body of data, existing resources are not ideal for resolving synovial joint formation. First, whole-limb atlases sample broadly across stages but capture relatively few cells per stage, so the IZ and its flanking populations, which are a small fraction of the LBM, are sparsely represented. Several of these atlases are derived from the forelimb, and several bracket the E11.5–E15.5 window in which joint progenitors are specified without covering it densely. Second, joint-focused datasets achieve resolution within the Gdf5 lineage or the IZ itself but, by design, omit the surrounding populations from which that lineage is recruited and against which its boundaries are set. Third, spatial datasets either predate IZ formation, lack single-cell resolution, or sample the limb only incidentally. Across all of these, cross-study comparability is limited by discordant nomenclature, differing sampling strategies (stage, anatomical domain, dissociation protocol), and the subtle transcriptional signatures of LBM cells as they diversify, which make it difficult to separate biologically meaningful populations from technical artifacts. Finally, the lack of a consensus cell-type taxonomy anchored to in situ spatial annotation makes it difficult to quantify the extent to which in vitro systems (micromass cultures, explants, and the recently described limb organoids) recapitulate normal development.

Here, we define the transcriptional identities of mouse hindlimb cells across synovial joint development by combining scRNA-seq with high-resolution spatial transcriptomics. We sampled the hindlimb densely from the stage of initial mesenchymal condensation (E11.5) through early knee cavitation (E15.5), profiling 228,703 cells across five daily stages and 47 spatial sections comprising 1,539,172 spots, at greater cell numbers per stage than existing atlases. This has allowed us to resolve the LBM-derived populations, their gene signatures, and their locations. We used this atlas to quantitatively assess the fidelity of limb explants to the in vivo hindlimb, and found that explant culture maintains the major lineages and the axial patterning of the developing limb for at least three days. We further used the atlas to identify transcriptional programs associated with mesenchymal condensation, and showed that one of their members, Sema3c, is required for chondrogenesis and for joint formation in explant culture. Moreover, we found that IZ cells carry a chondrogenic signature at E11.5 to E13.5 and a tenogenic signature at E14.5 to E15.5, and we uncovered previously unreported markers of the knee IZ, including Cntn4, Igfbp5, Ism1, Man1a, Pamr1, Prss23, Sema3c, Tafa2, Plxna4, Pdlim3, and Prkd1, of which Igfbp5 is required for knee formation in explants. Thus, this atlas provides a reference against which joint progenitor specification can be studied in vivo and assessed in vitro.

## RESULTS

### An integrated scRNA-seq and spatial transcriptomics atlas of the synovial joint

To define the cell states present during synovial joint formation and to anchor them to position in the limb, we profiled the developing mouse hindlimb at daily intervals from mesenchymal condensation (E11.5) to the onset of knee cavitation (E15.5) by scRNA-seq and spatial transcriptomics (Figure 1A). We used Sox9^-IRES-GFP^ embryos^39^, which express GFP from the native Sox9 locus. For scRNA-seq, we dissected the presumptive knee joint from the hindlimbs of 6 to 8 embryos per litter at E12.5 through E15.5. At E11.5, we collected the entire limb because the template is not yet discernable (Figure 1A). After enzymatic digestion, we depleted CD45^+^ and Ter119^+^ hematopoietic cells. At E11.5, E13.5, and E14.5, we loaded approximately 40,000 cells into a PIP-seq T20 reaction to roughly capture 20,000 cells per stage. To sample the initiation (E12.5) and the end (E15.5) of the interzone window more deeply, we loaded approximately 200,000 cells into a PIP-seq T100 reaction to capture roughly 100,000 cells per timepoint. For spatial transcriptomics, we placed sagittal and coronal sections of fresh-frozen hindlimbs onto 10mm-by-10mm Curio-seeker tiles, which sample the whole transcriptome at 10um resolution. After quality control, the atlas comprises 228,703 cells (16,011 at E11.5, 59,153 at E12.5, 24,484 at E13.5, 16,036 at E14.5, and 113,019 at E15.5) and 47 spatial sections comprising 1,539,172 spots (89,488 at E11.5; 318,996 at E12.5; 340,274 at E13.5; 520,120 at E14.5; 270,294 at E15.5) (Figure 1B).

**Figure 1.**
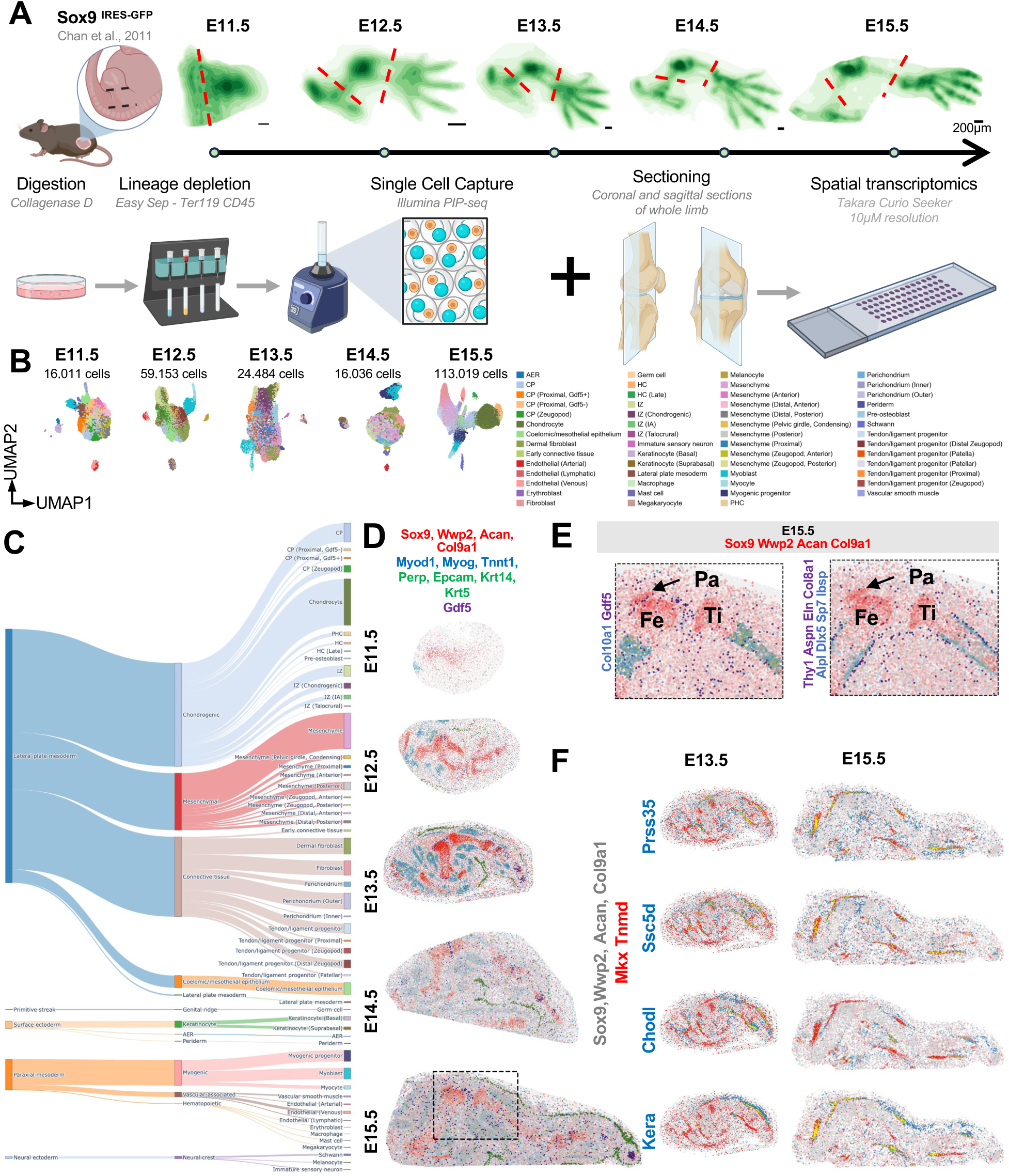
A single-cell and spatial transcriptomic atlas of the developing mouse hindlimb from E11.5 to E15.5. (A) Workflow. Hindlimbs of Sox9-IRES-GFP embryos [39] were collected daily from E11.5 to E15.5. Green, Sox9-GFP marking the cartilage template. Red dashed lines, the region dissected for scRNA-seq at each stage; the whole limb was taken at E11.5. For scRNA-seq, pooled limbs were digested with collagenase D, depleted of CD45+ and Ter119+ cells, and captured by PIP-seq. For spatial transcriptomics, coronal and sagittal sections of whole hindlimbs were placed on Curio Seeker tiles (10 um resolution). Scale bar, 200 um. (B) UMAP embeddings of the scRNA-seq data at each stage, colored by cell type, with the number of cells retained after quality control. Colors as in the legend at right. (C) Alluvial diagram relating embryonic lineage (left), broad cell class (center), and the 56 cell types (right) across all stages. Band width is proportional to cell number. (D) Representative spatial section at each stage. Red, cartilage template (weighted expression of Sox9, Wwp2, Acan, and Col9a1); blue, skeletal muscle (Myod1, Myog, Tnnt1); green, epidermis (Perp, Epcam, Krt14, Krt5); purple, Gdf5. (E) Perichondrial layers at E15.5 in the region boxed in (D). Red, cartilage template (Sox9, Wwp2, Acan, Col9a1). Left, blue, Col10a1 and Gdf5. Right, purple, outer perichondrial markers (Thy1, Aspn, Eln, Col8a1); blue, inner perichondrial markers (Alpl, Dlx5, Sp7, Ibsp). Arrow, Gdf5+ band between the patella and the distal femur. Pa, patella; Fe, femur; Ti, tibia. (F) Tendon and ligament markers at E13.5 and E15.5. Grey, cartilage template (Sox9, Wwp2, Acan, Col9a1); red, Mkx and Tnmd; blue, the indicated gene (Prss35, Ssc5d, Chodl, Kera).

We reasoned that integrating the five stages before clustering would blur the stage-specific variation that distinguishes LBM subpopulations, which differ from one another by subtle transcriptional signatures. We therefore clustered each stage separately and annotated clusters by determining where their marker genes are expressed in the limb, using both published in situ data and our own spatial sections (Figures S2 to S6). This yielded 11 broad classes and 56 subtypes across the five stages. The atlas covers the major lineages present of the limb: lateral plate mesoderm (mesenchyme, cartilage, and connective tissue), paraxial mesoderm (muscle, vasculature, and hematopoietic cells), surface ectoderm (epidermis and apical ectodermal ridge), and neural crest derivatives (Schwann cells and melanocytes) (Figure 1B, C). Subtypes defined by transcriptome alone localized to the expected anatomical positions in the spatial sections, which supports the notion that they are spatially distinct populations and not clustering artifacts (Figure 1D; Figures S2D to S6D). The spatial data resolve closely related tissues. For example, at E15.5 the developing patella is separated from the distal femur by a band of Gdf5 expression, and the perichondrium resolves into an outer layer marked by Thy1, Aspn, Eln, and Col8a1 and an inner layer marked by Alpl, Dlx5, Sp7, and Ibsp (Figure 1E).

Combining the two data types also yielded markers of tendon and ligament development that we had not anticipated. First, Prss35 labels developing tendons and ligaments from E13.5 to E15.5 (Figure 1F; Figures S4E, S6E). Prss35 is upregulated after medial collateral ligament injury in rat^40^,but to our knowledge it has not been localized to tendons or ligaments during development. PRSS35 also marks developing tendons and ligaments in the human hindlimb at 8 post-conception weeks (PCW) in two published spatial datasets (Figure S7)^29,41^, which suggests that Prss35 is a conserved marker of tendon and ligament development. Second, we find that Ssc5d is expressed in both the interzone and the developing tendons and ligaments. Ssc5d, Htra1^31^, and Aqp1 share a pattern that includes the tendons and the nascent intra-articular ligaments of the developing mouse and human hindlimb (Figure 1F; Figures S6E, S7). Scx binds to the basal promoter of Ssc5d and regulates its expression in the developing humeroulnar joint and forelimb tendons^42^. The presence of Ssc5d in tendon and ligaments of both the forelimb and hindlimb, in mouse and human, suggests that it is a part of a conserved tendon program. Third, Chodl, which is expressed in developing tendons^43^, is absent from the developing patellar ligament and nascent intra-articular ligament (Figure 1F). Finally, Kera is expressed along the anterior margin of the limb at E13.5 and E14.5, where it partially overlaps with Mkx expression in the extensor tendons (Figure S4E, 5E). By E15.5, Kera is restricted to developing tendons and ligaments in both the anterior and posterior compartments, including the patellar ligament (Figure 1F; Figure S6E). Using the combined patterns of Mkx, Tnmd, Chodl, Kera, and Ssc5d, we assigned the tendon and ligament progenitor clusters in the scRNA-seq data to patellar, proximal, and zeugopod locations (Figure 1C).

To test whether these transcriptional signatures are artifacts of our platform or dissection, we asked whether each cell type could be found in independent datasets. We classified seven published scRNA-seq datasets of the developing limb and joint^27,29,31,34,44–46^ using two methods: pySCN^47^, a random forest classifier trained on our annotated data, and Ingest^48,49^, which projects query cells into a PCA embedding of the reference and transfers labels by nearest-neighbor lookup. Except for rare populations outside the lateral plate mesoderm, every cell type we defined was detected in at least one other dataset by at least one method (Figure S1A, B).

We next compared the resolution of our lateral plate mesoderm annotations to those of two integrated atlases of mouse limb development^29,30^ by comparing pySCN classifier metrics for the lateral plate mesoderm populations of each atlas at each stage, restricting the published atlases to the stylopod and zeugopod where possible to match our dissection (Figure S1C). Three differences emerged. First, at E11.5 we defined 12 lateral plate mesoderm cell types, against 5 in each published atlas. The additional resolution comes from spatially distinct chondroprogenitor and mesenchymal progenitor populations; neither published atlas assigns a regional location to chondroprogenitors or an anterior or posterior identity to mesenchymal progenitors. Second, neither integrated atlas includes an annotated E14.5 stage, at which we defined 11 lateral plate mesoderm cell types. Third, at E15.5 we defined 15 cell types, against 7 and 10, because we separated several interzone and tendon and ligament progenitor populations that the published atlases label as “Interzone” or “Tenocyte precursors”. At E12.5 and E13.5, the number and identity of cell types were similar to those of Zhang et al. (12 versus 11 at E12.5; 15 versus 14 at E13.5). The Herpelinck et al. atlas had the highest median precision, recall, and F1 score at E11.5 and E13.5. We attribute this to its smaller number of cell types at those stages (5 and 11), since classifier metrics rise as classes are merged. Taken together, these comparisons indicate that the atlas recovers the cell types reported in prior studies, resolves the lateral plate mesoderm more finely at E11.5 and E15.5, and adds an annotated E14.5 stage that the integrated atlases lack.

### Limb explant culture maintains the lineages and axial patterning of the hindlimb

Fundamental insights into proximal-distal patterning and identification of key signaling pathways required for limb growth and maturation were enabled by the ability to manipulate limb development in ovo^50–53^, where populations can be maintained in situ following perturbation, and thus preserving the extracellular environment, including oxygen saturation, nutrient availability, and tissue architecture of accessory structures (sensory neurons, vasculature, skeletal muscle, and epithelial cells). Equivalent perturbations in the mouse require a transgenic line. In vitro alternatives, such as high-density micromass culture^54^ and embryonic stem cell-derived limb bud-like cultures^55^ are optimized for chondrogenesis and form many small nodules in place of discrete skeletal elements. Limb explant culture occupies the ground between these approaches. In the system described by Arostegui and Underhill, embryonic limbs are cultured partially submerged at an air-media interface for up to three days^56,57^ (Figure 2A). The method has since been adapted to rat limb buds^58^, and we recently showed that siRNA injected directly into the developing knee produces localized knockdown^59^. In our hands and others’, the cartilage template continues to grow over three days of culture and the knee region remains Sox9-negative, as judged by Sox9-GFP (Figure 2B)^59^. However, the extent to which explant culture preserves the transcriptional state of the limb has not, to our knowledge, been measured; prior characterization focused on autopod regeneration after WNT activation^60^ or the response to oxygen tension^61^. We therefore profiled the knee region of E12.5 explants by scRNA-seq, and whole explants by spatial transcriptomics, after 24 hours (D1) and 72 hours (D3) of culture (Figure 2B).

**Figure 2.**
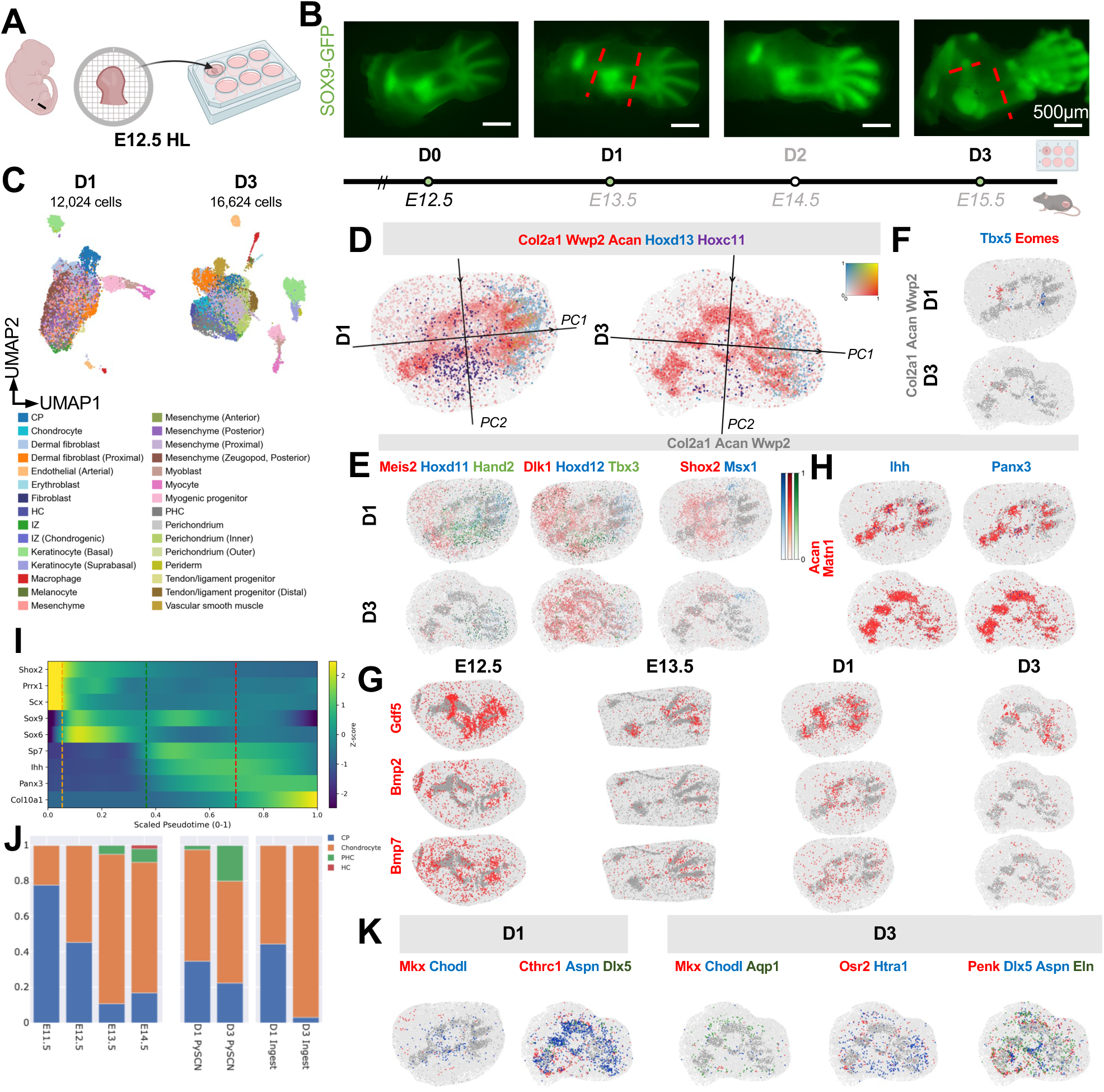
Limb explant culture maintains the lineages and axial patterning of the hindlimb but delays chondrocyte and connective tissue differentiation. (A) Explant culture. Hindlimbs were isolated at E12.5 and cultured at an air-media interface for up to three days. (B) Sox9-GFP in explants at D0 through D3. Samples were collected at 24 h (D1) and 72 h (D3) for scRNA-seq and spatial transcriptomics. Red dashed lines indicate dissection for scRNA-seq; whole explants were sectioned for spatial transcriptomics. The timeline gives the corresponding in vivo stages. Scale bar, 500 um (C) UMAP embeddings of explant scRNA-seq data at D1 (12,024 cells) and D3 (16,624 cells), colored by cell type. (D) Representative D1 and D3 sections with the principal axes of the spot coordinates: PC1 approximates the proximal-distal axis and PC2 the anterior-posterior axis. Red, cartilage (Col2a1, Wwp2, Acan); blue, Hoxd13; purple, Hoxc11. (E) Patterning genes along the proximal-distal and anterior-posterior axes at D1 and D3. Each panel shows three genes plotted in order: red (first), blue (second), green (third). Grey, cartilage template (Col2a1, Acan, Wwp2). (F) Tbx5 (blue) and Eomes (red) at D1 and D3. Tbx5 remains at the anterior margin and Eomes at the proximal end of the fourth digit throughout culture. Grey, cartilage template. (G) Interdigit mesenchyme in vivo (E12.5, E13.5) and ex vivo (D1, D3). Top, Gdf5 (red); middle, Bmp2 (red); bottom, Bmp7 (red). Grey, cartilage template. (H) Chondrocyte maturation markers at D1 and D3. Left, Ihh (blue); right, Panx3 (blue); both over Acan and Matn1 (red), with cartilage template (Col2a1, Acan, Wwp2) in grey. (I) Pseudotime of chondrocyte differentiation inferred with scFates from in vivo chondrocytes (E11.5 to E15.5). Heatmap, fitted expression of Shox2, Prrx1, Scx, Sox9, Sox5, Sox6, Sp7, Ihh, Panx3, and Col10a1 along scaled pseudotime. Dashed lines, boundaries between chondroprogenitor, chondrocyte, pre-hypertrophic, and hypertrophic states, set at inflection points of the marker curves. (J) Proportions of chondrocyte states by stage in vivo (E11.5 to E15.5) and in explants (D1, D3) as classified by pySCN and by Ingest. CP, chondroprogenitor; PHC, pre-hypertrophic chondrocyte; HC, hypertrophic chondrocyte. K) Connective tissue populations at D1 and D3. D1: Mkx (red) and Chodl (blue); Cthrc1 (red), Aspn (blue), and Dlx5 (green). D3: Mkx (red), Chodl (blue), and Aqp1 (green); Osr2 (red) and Htra1 (blue); Penk (red), Dlx5 and Aspn (blue), and Eln (green). Grey, cartilage template.

After quality control, we retained 12,024 cells at D1 and 16,624 cells at D3, from two replicate cultures per timepoint (Figure 2C). Explants retained the lateral plate mesoderm, paraxial mesoderm, and surface ectoderm lineages at both timepoints (10,798, 888, and 338 cells at D1; 13,134, 1,894, and 1,555 cells at D3), as well as hematopoietic cells (41 macrophages at D1; 123 macrophages and 49 erythroblasts at D3). Neural crest-derived melanocytes were detected at D3 (41 cells) but not at D1. We annotated 18 cell types at D1 and 27 at D3, and the D3 explant contained chondrocyte, pre-hypertrophic chondrocyte, hypertrophic chondrocyte, inner and outer perichondrial, tendon and ligament progenitor, and interzone populations that match in vivo annotations (Figure 2C; Figure S8A, B). These observations are consistent with a recent report that explants sustain the major lineages of the limb for at least three days^61^.

We then asked whether explant culture preserves patterning along the limb axes. Using our spatial transcriptomics data, we examined the genes that vary across the proximal-distal axis and anterior-posterior axis. For each section, principal component analysis (PCA) of the spot coordinates gives a first component (PC1) that aligns with the proximal-distal axis and a second (PC2) that approximates the anterior-posterior axis (Figure 2D). We projected spots onto each axis and tested whether either the expression level or the fraction of expressing spots changed along it. For in vivo stages, we retained genes that changed in the same direction in at least two sections per stage. Along PC1, the distal Hox genes Hoxa13, Hoxd10, Hoxd11, Hoxd12, and Hoxd13, and Msx1, increased toward the distal end in every in vivo stage and explant timepoint (Figure 2E; Figure S8C). Meis1, Meis2, and Shox2 decreased along PC1 in vivo and ex vivo, consistent with their proximal restriction (Figure S8C). Sall1 and Zic3 localized to the zeugopod in explants, as they do at E12.5 and E13.5 in vivo (Figures S3E, S4F). Along PC2, Kera marked the anterior margin at E13.5 and E14.5 (Figures S4E, S5D) and throughout culture (Figure 2E), and Hoxc11 marked the posterior margin in vivo and ex vivo (Figure 2E; Figures S3E, S4F). Tbx5, which is usually associated with the forelimb, was expressed along the anterior margin of the knee interzone at E12.5 and E13.5 and remained there in explants (Figure 2F; Figures S3E, S4F).

However, we found that explant cultures diverged from in vivo development in the autopod. In vivo, the digit rays are separated at E12.5 and E13.5 by interdigit mesenchyme that expresses Gdf5, Bmp2, and Bmp7. After one day in culture the digits were less defined. Gdf5 expression persisted between the digits, but Bmp2 and Bmp7 were not detected in the interdigit mesenchyme at D1 or D3 (Figure 2G). Eomes expression at the proximal end of the fourth digit was maintained throughout culture despite the malformation of the autopod (Figure 2F).

### Chondrocyte hypertrophy and connective tissue differentiation is stunted in limb explant culture

The skeletal malformations in the autopod lead us to ask to what extent chondrocyte differentiation proceeds in explant culture. A Gdf5^+^ interzone population was present in both the scRNA-seq and spatial transcriptomics data throughout culture (Figure 2G). At D1, this population also expressed high levels of Sox9 and Col2a1, consistent with the E12.5 and E13.5 interzone in vivo (Figure S3B, S4B, S8A). Outside of the interzone, the Sox9^+^ Col2a1^+^ cartilage template expressed Col9a1^+^ and Wwp2^+^ throughout the culture period. Acan and Matn1 displayed delayed onset: at D1 they were restricted to the diaphysis and base of the metatarsals, and by D3, expression was present throughout the Sox9^+^ cartilage template. The pre-hypertrophic markers Sp7 and Ihh were present in the diaphysis by D1, and Panx3 by Day 3 (Figure 2H). To place these observations on a common scale, we aligned in vivo chondrocytes from E11.5 to E15.5 along a pseudotime of chondrocyte differentiation using scFates^62^ and defined chondroprogenitor, chondrocyte, pre-hypertrophic chondrocyte (PHC), and hypertrophic chondrocyte (HC) states from the inflection points of Sox9, Sox5, Sox6, Sp7, Ihh, Panx3, and Col10a1 (Figure 2I). We then classified explant chondrocytes against these states with pySCN and with Ingest. The two methods disagreed on the most advanced state present: pySCN assigned explant cells to chondroprogenitor, chondrocyte, and PHC states, whereas Ingest assigned none to the PHC state. The methods agreed on two points: the proportion of chondroprogenitors fell from D1 to D3, and no explant cell was classified as a hypertrophic chondrocyte (Figure 2J). Chondrocytes therefore mature in culture but do not complete hypertrophic differentiation within three days.

Explant culture also contained connective tissue populations whose differentiation lagged the in vivo sequence. At D1, we identified two early populations. One expressed the tendon and ligament progenitor markers Mkx and Chodl^63^ together with the posterior markers Hand2 and Hoxc11, and the other expressed the perichondrial markers Dlx5, Aspn, and Cthrc1 (Figure 2K). By D3, the tendon progenitor population expressed Mkx, Scx, Tnmd, and Chodl, and a second Osr2^+^ Htra1^+^ population with low Chodl resembled the ligament progenitors of the E15.5 hindlimb (Figure S6B). In vivo, Mkx^+^ Tnmd^+^ tendon and ligament condensations are visible by E13.5. In explants, no such structures were visible at D1, but by D3 Mkx, Chodl, Aqp1, Scx, Osr2, and Htra1 localized adjacent to the cartilage template (Figure 2K). At D3 we also found two Dlx5^+^ perichondrial populations. The first expressed higher levels of Sox9 and Cthrc1, as do the inner cambial perichondrial cells at E15.5 in vivo (Figure 1E; Figure S6B), but lacked the Ibsp and Sp7 expression of their in vivo counterparts. The second expressed higher levels of Aspn, Eln, and Penk, and Dlx5, Aspn, Eln, and Penk expression directly bordered the cartilage template at D3 (Figure 2K), which suggests that this population corresponds to the outer fibrous perichondrium.

Taken together, these analyses demonstrate that limb explant culture maintains the major cell populations, the axial patterning genes, and transcriptional programs observed in the E12.5 to E15.5 hindlimb, with the exception of interdigit patterning, and that chondrogenic and connective tissue differentiation lag behind their in vivo counterparts. The system is therefore suited to functional studies of the knee over this window, with the delay in differentiation to be kept in mind when a phenotype depends on hypertrophy or on mature tendon.

### Sema3c is co-expressed with Sox9 and Gdf5 during condensation and is required for chondrogenesis and joint formation in culture

Having defined the cell states of the developing limb both in vivo and in explant culture, we next asked how the transcriptome changes during mesenchymal condensation and early chondrogenesis.

At E11.5, the characteristic Y-shaped pre-cartilage condensation (PCC) of the hindlimb is subtly defined by Sox9 and Col2a1 expression. In line with previous reports, we found that the interzone marker Gdf5 is already expressed in the proximal limb at this stage^22,64^, and in our spatial transcriptomic sections, its domain overlaps with the developing femoral template (Figure 3D). Gdf5 enhances chondrogenesis during mesenchymal condensation, and its overexpression of Gdf5 increased the length and width of skeletal elements^18,64^. Consistent with the relationship between Sox9 and Gdf5, reanalysis of published RNA-seq data^65^ demonstrated a decrease in Gdf5 expression in mice where Sox9 has been knocked out in Prrx1-Cre expressing cells compared to controls (Figure S9A). We therefore sought other genes that have dynamic expression during mesenchymal condensation and chondrogenesis and might influence Sox9 or Gdf5 expression.

**Figure 3.**
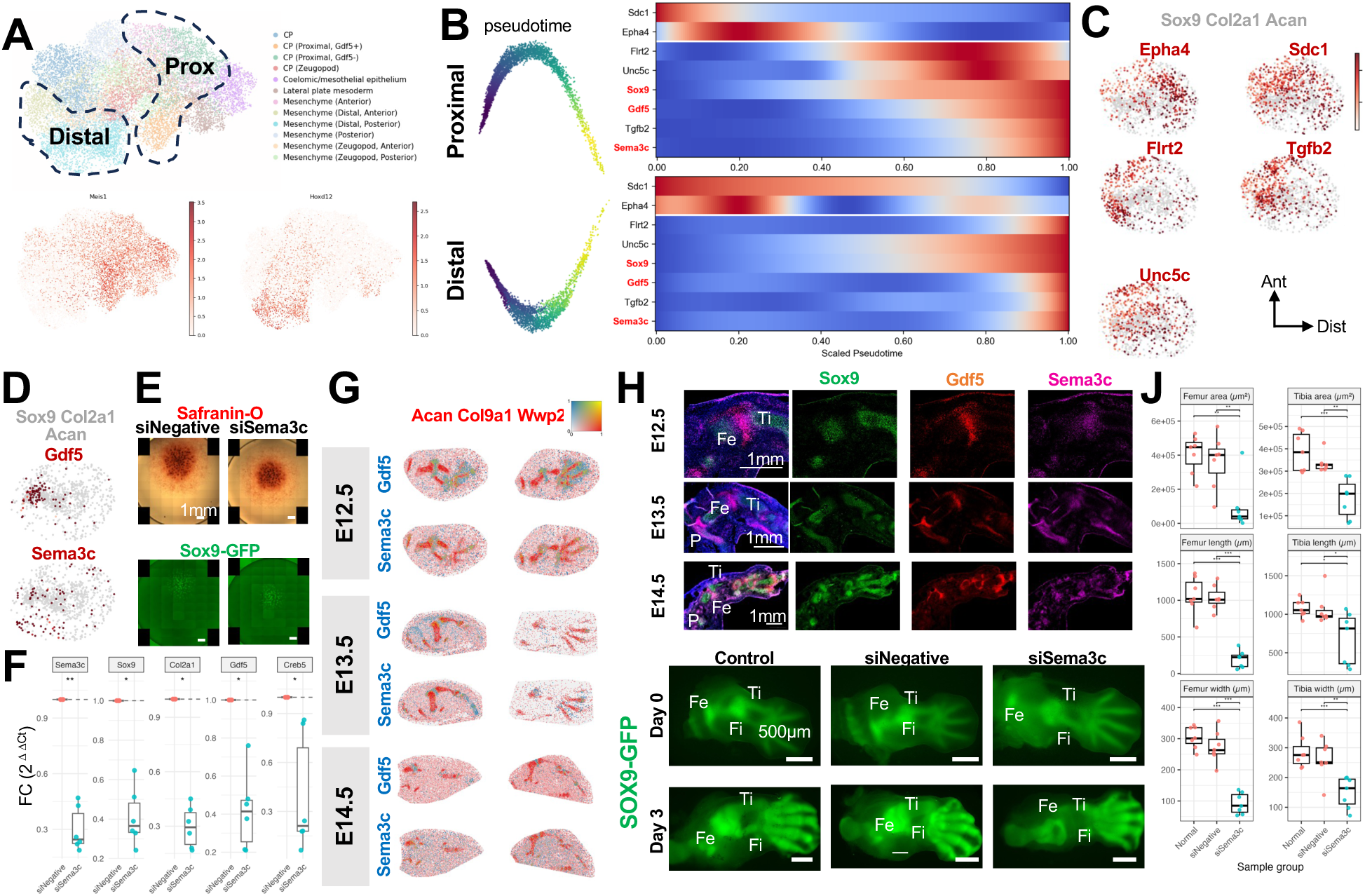
Sema3c is co-expressed with Sox9 and Gdf5 during condensation and is required for chondrogenesis and joint formation in culture. A) UMAP of E11.5 Prrx1+ lateral plate mesoderm colored by cell type, with the proximal and distal subsets used for trajectory inference outlined. Below, Meis1 (proximal) and Hoxd13 (distal) expression. (B) Left, diffusion map embeddings of the proximal and distal subsets colored by scFates pseudotime. Right, fitted expression of Sdc1, Epha4, Flrt2, Unc5c, Sox9, Gdf5, Tgfb2, and Sema3c along scaled pseudotime for each trajectory. Genes in red are referred to in the text. (C) Spatial expression at E11.5 of Epha4, Sdc1, Flrt2, Tgfb2, and Unc5c (red) over the pre-cartilage condensation (Sox9, Col2a1, Acan; grey). Axes: Prox, proximal; Dist, distal; Ant, anterior; Post, posterior. (D) E11.5 section showing Gdf5 (top, red) and Sema3c (bottom, red) over the condensation (Sox9, Col2a1, Acan; grey). Sema3c+ Gdf5+ cells lie in the proximal femoral template. (E) Safranin-O staining (top) and Sox9-GFP (bottom) of E11.5 hindlimb micromass cultures 48 h after transfection with scrambled (siNegative) or Sema3c siRNA. Scale bar, 1 mm. (F) RT-qPCR of Sema3c, Sox9, Col2a1, Gdf5, and Creb5 in micromass cultures after Sema3c knockdown, as fold change (2^-ddCt) relative to siNegative. n = 6 cultures per condition; three technical replicates per gene. Welch’s two-sample t-test on dCt values with Bonferroni correction across genes., adjusted p < 0.05;, adjusted p < 0.01; **, adjusted p < 0.001. (G) Spatial expression of Gdf5 and Sema3c (blue) at E12.5, E13.5, and E14.5 over the cartilage template (Acan, Col9a1, Wwp2; red).| (H) RNAscope for Sox9 (green), Gdf5 (red), and Sema3c (magenta) in the knee at E12.5, E13.5, and E14.5; merged image with DAPI at left. Fe, femur; Ti, tibia; P, patella. Scale bar, 1 mm. (I) Sox9-GFP in E12.5 hindlimb explants at D0 and D3 after no injection (Control), scrambled siRNA (siNegative), or Sema3c siRNA injected into the knee (n = 7 for each condition). Fe, femur; Ti, tibia; Fi, fibula. Scale bar, 500 um. (J) Femoral and tibial area, length, and width at D3. One-way ANOVA followed by Tukey’s HSD; siNegative did not differ from Control for any measure.

To examine the transcriptomic changes associated with mesenchymal condensation and early chondrogenesis, we applied scFates to reconstruct pseudo-temporal trajectories of the proximal and distal E11.5 limb bud mesenchyme. Since chondrogenesis progresses asynchronously across the proximo-distal axis, we generated independent trajectories for each region (Figure 3a). We restricted the analysis to Prrx1-expressing, which marks the lateral plate mesoderm lineage that gives rise to osteochondral progenitors^7,66^. For each subset, we built a diffusion map from the top 100 and bottom 100 genes correlated with Sox9 by biweight midcorrelation^67^ and inferred a trajectory on it. We then grouped genes by their expression profile along pseudotime and examined the groups whose expression changed (Figure 3B; Figure S9C, D). Some genes peaked at different points in the two trajectories. We reasoned that the distal trajectory captures early condensation, since the distal mesenchyme is less differentiated, whereas the proximal trajectory captures late condensation and early chondrogenesis (Figure S9B). On this reading, genes that peak early in the proximal trajectory and late in the distal trajectory are condensation genes that decline as chondrogenesis begins. This group included Slit2, Col1a1, Col1a2, Col3a1, Igf1, Pth1r, Alcam, Dlk1, Ednra, and Lpar1 (Figure S9C, D).

Many genes showed consistent expression patterns in both trajectories, which suggests a single program running from condensation through chondrogenesis, and this group included established regulators of both processes. Sdc1 and Epha4^68,69^ peaked early in both trajectories. In the spatial transcriptomics data, Sdc1 surrounded the pre-cartilage condensation (PCC), while Epha4 was expressed along the distal edge of the PCC (Figure 3C). Tgfb2, Flrt2, and Unc5c peaked later in both trajectories^70^; Tgfb2 and Flrt2 were expressed in the proximal limb, including the PCC, while Unc5c is expressed throughout the template (Figure 3C). Three semaphorin ligands (Sema6a, Sema3c, and Sema3d) showed similar pseudotime profiles to that of Gdf5 (Figure S9B, C). Sema3c and Sema3d have been implicated in condensation: both are reduced in Sox9-null limbs, and Sema3c knockdown in high-density micromass culture reduces Alcian blue staining^65^.The effect of Sema3c on specific chondrogenic genes has not, to our knowledge, been measured. Sema3c expression was sparse at E11.5. In the scRNA-seq data, 89 of 16,011 cells had detectable Sox9, Gdf5, and Sema3c transcripts, and 57 of these were in the Gdf5+ proximal chondroprogenitor cluster (Figure S9E). In the spatial sections, the co-expressing cells lay in the proximal femoral template (Figure 3D). We therefore tested whether Sema3c loss affects these genes. We transfected E11.5 hindlimb mesenchyme with Sema3c siRNA or a scrambled control at the time of micromass seeding and measured expression after 48 hours. Sema3c knockdown reduced Sox9, Col2a1, and Gdf5 expression (Figure 3F; n = 6 cultures per condition, Welch’s t-test on delta-Ct with Bonferroni correction). Creb5, an interzone transcription factor required for joint formation^71,72^, was also reduced, which suggests that Sema3c may act in the early interzone in addition to the condensation.

Because Sema3c knockdown reduced Gdf5 and Creb5, we asked whether Sema3c is expressed in the interzone at later stages (Figure 3G, H). At E12.5 and E13.5, Sema3c was expressed at high levels in the interzone and the perichondrium. By E14.5, its expression had become diffuse throughout the mesenchyme, which suggests that Sema3c acts during interzone formation but is not required for its maintenance. SEMA3C was likewise expressed in the developing human knee at 6.2 to 7.1 PCW, roughly equivalent to E12.5 to E13.5 in mouse but was no longer restricted to the interzone at 8.1 PCW, roughly E15.5 to E16.5 (Figure S9E)^29^.

We then tested the requirement for Sema3c in the early interzone by injecting Sema3c siRNA into the developing knee of E12.5 explants and culturing them for three days. Sema3c knockdown reduced femoral and tibial area, length, and width relative to uninjected and scrambled-siRNA controls, and no joint was visible after three days (Figure 3I, J; n =7 limbs per condition, one-way ANOVA with Tukey’s HSD; the scrambled control did not differ from the uninjected control for any measure). Taken together, these results indicate that Sema3c promotes chondrogenesis and that its loss impairs joint formation in culture.

The expression of Sema3c, enriched in the early interzone and diffuse by E14.5, raised the broader question of how the interzone itself changes over developmental time. We therefore examined interzone cell states from E12.5 to E15.5.

### Interzone cells carry a chondrogenic signature early and a tenogenic signature late

Lineage tracing has shown that every component of the synovial joint descends from cells that expressed Gdf5, and that the interzone is fed by a continuous influx of Gdf5-expressing cells from the surrounding mesenchyme^29,31,34,44–46^. Cells that express Gdf5 early preferentially populate the epiphysis, whereas late expressing Gdf5 cells contribute to the articular cartilage and fibrous components^22^. Gdf5-lineage cells rapidly emigrate from the interzone and downregulate Gdf5 expression^22^, such that Gdf5-lineage cells and actively Gdf5-expressing cells overlap, but are not completely co-localized. We first asked whether our data reflect this transience. If cells that have expressed Gdf5 persisted in an interzone transcriptional state, then a dataset enriched for the Gdf5 lineage should contain a higher fraction of cells classified as interzone than a dataset that is not enriched. We classified published scRNA-seq datasets of the limb and knee^29,31,34,44–46^ from E12.5 to E15.5 with our stage-specific pySCN classifiers and counted the fraction of limb mesenchyme classified as interzone (Figure S10). At E12.5, the fractions were similar (16.2% in the Gdf5-lineage dataset versus 14.8% and 13.7%). At E13.5 and E14.5, the Gdf5-lineage-enriched datasets^31^ had a lower fraction of interzone cells than the non-enriched datasets (7.1% versus 10.6% to 34.2% at E13.5; 1.5% versus 8.0% and 10.1% at E14.5). Interestingly, at E15.5, the Gdf5-lineage and Prrx1-Cre datasets had a large proportion of cells classified as intra-articular interzone (28.7% and 25.3% versus 8.8%) (Figure S10C). These fractions do not increase with lineage enrichment at E12.5 to E14.5, which is consistent with interzone cells being transient. Given this transience, we asked whether interzone cells share a transcriptional profile across stages, or whether they are united only by Gdf5 expression.

We began with the transition from pre-interzone to interzone. We trained a pySCN classifier on the E12.5 data, where the interzone is morphologically distinct, and applied it to the E11.5 data, where it is not. Some Gdf5+ chondroprogenitors at E11.5 were classified as E12.5 interzone, but most were classified as E12.5 chondrocytes (Figure S11). We observed the same result in two independent E11.5 datasets, from C57BL/6J embryos^27^ and from Prrx1-Cre lineage-traced embryos^73^ (Figure S11).

For each of E12.5, E13.5, E14.5, and E15.5, we split the cells into training and held-out sets and trained a classifier on the training set, which gave four stage-specific classifiers. We applied each classifier to the held-out cells of all four stages, so that every cell received four labels. We kept cells labeled interzone by at least one classifier and discarded cells labeled interzone by non-consecutive stages, because a cell that resembles the E12.5 and E14.5 interzone but not the E13.5 interzone is unlikely to reflect a biological trajectory. This left ten categories: four single-stage (E12.5, E13.5, E14.5, E15.5), three two-stage (E12.5 to E13.5, E13.5 to E14.5, E14.5 to E15.5), two three-stage (E12.5 to E14.5, E13.5 to E15.5), and one spanning all four stages. For each stage, we found the genes differentially expressed among the ten categories and prioritized those that replicated in at least two datasets (Figure 4A). This design uses the signature of each stage on its own terms, separates transient from sustained interzone identity, and favors genes that reproduce across datasets.

**Figure 4.**
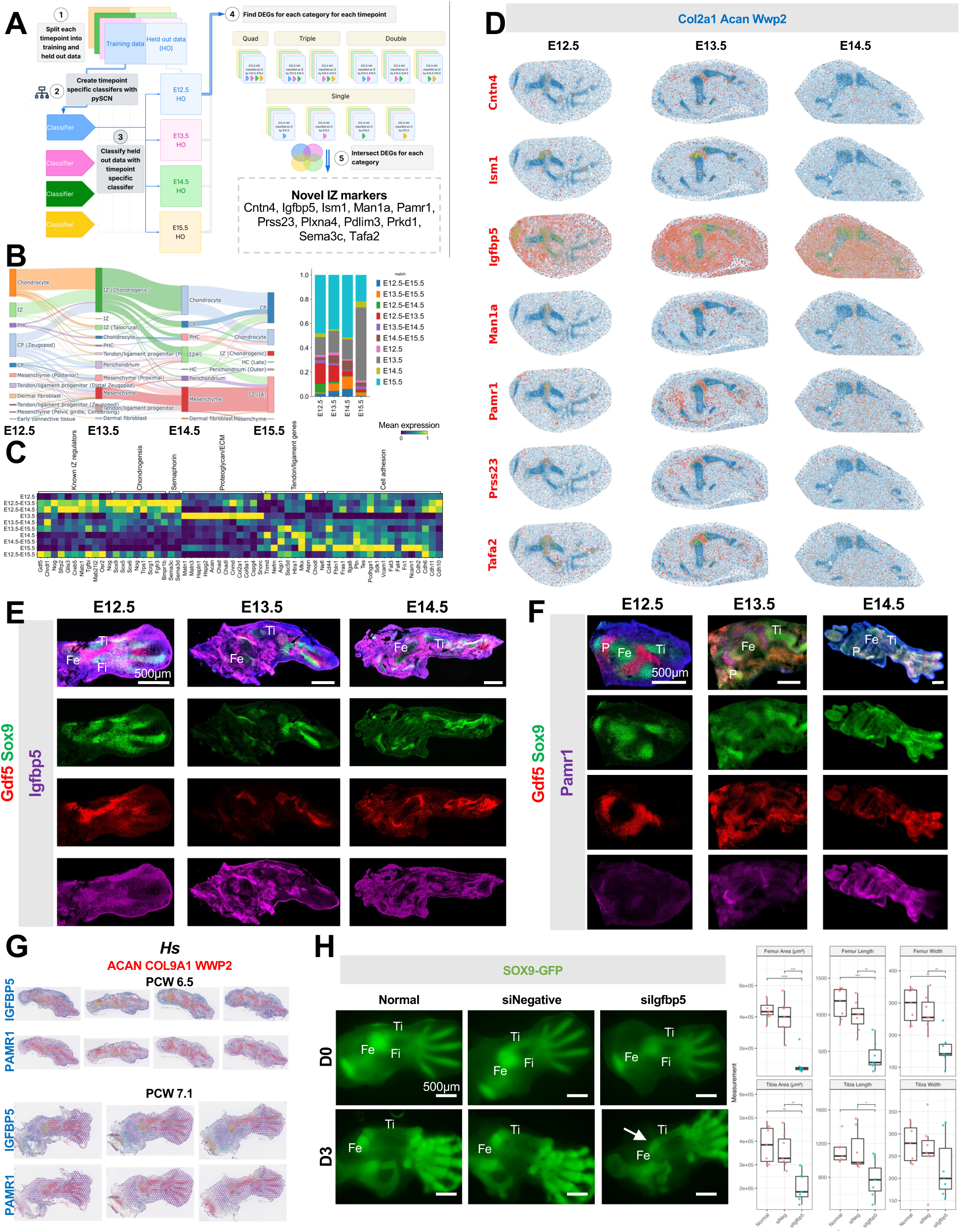
Interzone identity shifts from chondrogenic to tenogenic across stages, and a core signature includes previously unreported markers required for joint formation. (A) Cross-stage interzone analysis. (1) Cells at each stage were split into training and held-out sets. (2) A pySCN classifier was trained for each stage. (3) Each classifier was applied to the held-out cells of every stage. (4) Cells labeled interzone by at least one classifier were grouped into ten categories by which consecutive classifiers labeled them (single, double, triple, quad), and differentially expressed genes were found for each category at each stage. (5) Genes replicating across datasets were intersected. The box lists the interzone markers that have not, to our knowledge, been reported previously. (B) Left, alluvial diagram of cells labeled interzone by at least one classifier, showing the label each stage-specific classifier assigned. Right, fraction of each stage’s cells in each of the ten categories. (C) Mean expression, per category, of selected genes grouped as known interzone regulators, chondrogenic, semaphorin, perichondrial and extracellular matrix, tendon and ligament, and cell adhesion genes. Rows are the ten categories; columns are genes. (D) Spatial expression of Cntn4, Igfbp5, Ism1, Man1a, Pamr1, Prss23, and Tafa2 (red) at E12.5, E13.5, and E14.5 over the cartilage template (Col2a1, Acan, Wwp2; blue). Fe, femur; Ti, tibia; Fi, fibula; Pe, pelvis. (E, F) RNAscope for Gdf5 (red), Sox9 (green), and Igfbp5 (E) or Pamr1 (F) (magenta) in the knee at E12.5, E13.5, and E14.5; merged image with DAPI at top. Fe, femur; Ti, tibia; P, patella. Scale bar, 500 um. (G) IGFBP5 and PAMR1 (blue) in published spatial transcriptomics sections of the human hindlimb at 6.5 and 7.1 PCW [29] over the cartilage template (ACAN, COL9A1, WWP2; red). (H) Left, Sox9-GFP in E12.5 explants at D0 and D3 after no injection (Normal), scrambled siRNA (siNegative), or Igfbp5 siRNA (n = 7 per condition). Arrow, fused tibiofemoral joint. Fe, femur; Ti, tibia; Fi, fibula. Scale bar, 500 um. Right, femoral and tibial area, length, and width at D3. One-way ANOVA followed by Tukey’s HSD; siNegative did not differ from Normal for any measure.

The interzone was heterogeneous across time. Many cells were labeled interzone by only one or two classifiers (Figure 4B; Figure S12A). Cells labeled interzone by the early classifiers had the highest expression of chondrogenic genes, in line with the E11.5 result (Figure 4C). Likewise, a published Col2a1-lineage-traced dataset at E13.5^45^ contains Gdf5^+^ cells that resemble the E12.5 and E13.5 interzone, and these cells express high levels of Col2a1 (Figure S12B-D). In contrast, cells labeled interzone only by the late classifiers were enriched for tenogenic genes such as Tnmd, Chodl, and Aspn (Figure 4C).

Because the cell type labels themselves could impose this pattern, we repeated the analysis without them, using two methods to find genes co-expressed with Gdf5 at each stage. First, we computed a gene-by-gene biweight midcorrelation matrix^67^ and grouped genes into modules by average-linkage clustering. Second, we treated genes as observations and cells as features, found the nearest neighbors of Gdf5 in that space, and extracted gene programs by non-negative matrix factorization (NMF)^74^. Both methods placed Gdf5 in a module of chondrogenic genes at E12.5 and E13.5 (Figure 4D, E; Figure S13). Sox5, Sox6, Sox9, Col2a1, Col9a3, Cthrc1, Mia, Papss2, and Wwp2 were among the 50 genes most similar to Gdf5 by both methods at these stages. At E14.5 and E15.5, Gdf5 fell outside every correlation module and every NMF program, and its correlation with chondrogenic genes declined (Sox9: 0.40 at E12.5, 0.46 at E13.5, 0.23 at E14.5, below threshold at E15.5; Col2a1: 0.38, 0.28, below threshold, 0.15; Figure 4D). The chondrogenic program and Gdf5 therefore lose their coordination after E13.5.

Taken together, these analyses suggest that interzone cells, while they express Gdf5, already express genes of the lineages they will later join, a pattern reminiscent of lineage priming in hematopoietic stem cells. Early interzone cells (E11.5 to E13.5) resemble chondroprogenitors, and late interzone cells (E14.5 to E15.5) resemble other lateral plate mesoderm derivatives such as tendon and ligament progenitors.

### A core interzone signature shared across stages includes previously unreported markers

Despite this heterogeneity, a set of genes was enriched in cells classified as interzone by several classifiers, and we take these as a core interzone signature (Figure 4A, C). The interzone expresses a composite program: chondrogenic genes shared with the template, mesenchymal genes shared with the surrounding tissue, and genes specific to the interzone. The specific set includes known interzone regulators, among them the Wnt modulators Dkk3 and Sfrp2^75^ and the transcription factors Nfatc1^76^, Creb5^71^ and Osr2^77^(Figure 4C). It also includes genes that have not, to our knowledge, been reported as interzone markers: Cntn4, Igfbp5, Ism1, Man1a, Pamr1, Plxna4, Pdlim3, Prkd1, Prss23, Sema3c, and Tafa2. Each was enriched in the interzone at two or more stages in the spatial sections, with the most prominent localization to the interzone at E12.5 and E13.5 (Figure 4F; Figures S14, S15). Human homologs of eight of them (CNTN4, IGFBP5, ISM1, PAMR1, PLXNA4, PRSS23, PRKD1, and TAFA2, also known as FAM19A2) were expressed in the developing human knee at 6.2 to 7.1 PCW in published spatial data^29,41^ (Figure 4I; Figure S16). We confirmed the interzone localization of Igfbp5 and Pamr1 by RNAscope at E12.5, E13.5, and E14.5 (Figure 4G, H).

Among these genes, Igfbp5 had prior evidence of a role in joint biology. Its expression is elevated in Kashin-Beck disease, an osteochondropathy marked by enlarged joints and shortened digits^78^ and reduced in chondrocytes from the femoral condyles and tibial plateaus of osteoarthritis patients ^79,80^. Its role in joint development has not, to our knowledge, been tested. Igfbp5 is also expressed in early muscle condensations and in the interdigit mesenchyme^81,82^ (Figure 4F, G, I), so we delivered Igfbp5 siRNA directly into the tibiofemoral interzone of E12.5 explants and cultured them for three days. Igfbp5 knockdown produced a fused knee, and reduced femoral area, length, and width and tibial area and length relative to both controls (Figure 4J; n = 7 limbs per condition, one-way ANOVA with Tukey’s HSD; the scrambled control did not differ from the uninjected control). Igfbp5 is therefore required for tibiofemoral joint formation in explant culture.

## Discussion

A long-standing goal of limb biology is to explain how the multipotent limb bud mesenchyme partitions itself into cartilage, joint, tendon, and the other connective tissues, and in particular how the interzone recruits some neighbors and excludes others. Progress on this question has been limited by the available references: whole-limb atlases that sample the interzone sparsely, joint-focused datasets that omit the surrounding mesenchyme, and spatial datasets that predate interzone formation or lack cellular resolution. Here, we describe an atlas of the mouse hindlimb from E11.5 to E15.5 that pairs 228,703 single-cell transcriptomes with 28 spatial sections, annotates 56 cell types by their position in the tissue, and extends the same profiling to limb explant culture.

Several findings from this atlas have implications for the study of joint formation. First, explant culture preserves the lineages, the axial patterning genes, and the transcriptional programs of the hindlimb for three days, with two departures: interdigit Bmp2 and Bmp7 expression is lost by D1, and chondrocyte hypertrophy and tendon maturation lag their in vivo schedule. Explant culture is therefore a faithful system for perturbations of the knee between E12.5 and E15.5, and we used it to test two genes in days; a conditional allele would have taken months. Second, Sema3c, which we found co-expressed with Sox9 and Gdf5 in the condensing femoral template, is required in micromass culture for Sox9, Gdf5, and Creb5 expression and in explant culture for joint formation. Sema3c is enriched in the interzone at E12.5 and E13.5 and diffuse by E14.5, which suggests that chondrogenic processes continue inside the early interzone. Finally, interzone cells carry a chondrogenic signature early and a tenogenic signature late, and a core set of genes, including eleven that had not been reported as interzone markers, is shared across stages.

The chondrogenic signature of early interzone cells admits two interpretations. One is biological: Gdf5-expressing cells are biased toward a lineage before or at the time they turn on Gdf5, and do not acquire fate only after they arrive. The other is technical: the stage-specific classifiers could assign a chondrogenic label to early interzone cells if the early references contained mostly undifferentiated chondrocytes and the late references mostly pre-hypertrophic and hypertrophic chondrocytes, so that early interzone cells matched the nearest available class. Two observations argue against the technical reading. First, the annotation-free analyses, biweight midcorrelation and NMF, place Gdf5 in a chondrogenic module at E12.5 and E13.5 and nowhere at E14.5 and E15.5, without reference to any label. Second, pseudotime analysis shows that Sox9, Sox5, and Sox6 peak in chondrocytes and decline at the pre-hypertrophic transition, consistent with published reports, so the early references are not depleted of differentiated states. We therefore favor the biological reading.

How does this reading fit the prevailing model, in which Gdf5+ interzone cells arise by de-differentiation of Sox9+ Col2a1+ chondroprogenitors^5,83^? Col2a1-CreER labeling shows that Col2a1 is induced early, around E11.5, in the prospective joint region^84^, yet the E13.5 interzone contains both Col2a1-CreER+ and Col2a1-CreER-cells when labeled at E12.5 ^45^. Our data show that interzone cells at E12.5 and E13.5 express Col2a1 at levels comparable to Gdf5-chondrocytes, though below those of pre-hypertrophic chondrocytes, and that the Gdf5 lineage does not persist in an interzone state. We propose that the decline of Col2a1 at the site of the new interzone is not solely a matter of resident Col2a1+ Gdf5+ cells de-differentiating. Some Gdf5+ chondrogenic cells are displaced from the interzone and contribute to the skeletal elements, while newly arriving Gdf5-expressing cells replenish the interzone. This extends the Gdf5 influx model^22^, in which continuous immigration of Gdf5-expressing cells is required for joint formation, by adding relocation as a second contributor to the apparent loss of chondrogenic markers.

The core interzone genes point to a mechanism for one of the interzone’s defining properties, its exclusion of blood vessels. Several of the genes regulate vascularization: Ism1^86^ and Igfbp5^87^ are anti-angiogenic, and Plxna4 is pro-angiogenic in endothelial cells^88^; others, Man1a^89^ and Pamr1^90^, are predicted to bind calcium. The vascular regulators are of particular interest given that the interzone and the articular cartilage it produces are avascular. Sema3c, which the cross-stage analysis also recovered, causes vascular regression in several developmental contexts by inducing cytoskeletal collapse in endothelial cells through Plxnd1^91–93^, and we detect Plxnd1 in embryonic endothelial cells. Sema3c also binds Plxna4, which we found in the interzone but not in endothelium during joint development, which suggests a cell-type-specific role for Plxna4 in the interzone distinct from its pro-angiogenic role elsewhere. Sema3c, Ism1, and Igfbp5 are secreted and act directly on endothelial cells to cause repulsion or apoptosis, which would create a diffusible anti-angiogenic environment. Taken together, these observations suggest that the interzone uses several complementary mechanisms to establish and keep an avascular microenvironment. How the same tissue excludes endothelial cells while recruiting joint progenitors remains an open question.

Several limitations of this study warrant consideration. The developing limb contains multiple interzone populations corresponding to different joints. While these populations are spatially distinct, it remains unclear whether joint-specific mechanisms may confound our analysis. Although we attempted to isolate the developing tibiofemoral joint based on Sox9 expression, other interzone populations may contaminate our scRNA-seq data. Additionally, while we demonstrate that Sema3c and Igfbp5 knockdown resulted in joint deformities in explant culture, we cannot distinguish direct transcriptional effects from indirect consequences of failed condensation or vascular dysregulation. Finally, the function of Plxna4 in the interzone and the potential role of calcium signaling remain open questions for future investigation.

We acknowledge that this study is limited in several ways, which suggest avenues for future work. First, the developing limb contains several interzones, and although we dissected toward the tibiofemoral joint using Sox9-GFP, cells from other joints may be present in the scRNA-seq data. The spatial sections mitigate this, because every annotation was checked against position, and at E13.5 the talocrural interzone separated from the knee interzone as its own cluster (571 cells). Second, the Sema3c and Igfbp5 knockdowns produce joint defects in explant culture, but we cannot yet separate direct transcriptional effects in the interzone from indirect consequences of impaired condensation or vascular dysregulation. Profiling knockdown explants by scRNA-seq, which the explant atlas now makes interpretable, is the direct test. Third, explant culture delays hypertrophy and tendon maturation, so phenotypes that depend on either should be read with that delay in mind, or tested in vivo. Finally, the functions of Plxna4 and of the calcium-binding genes in the interzone are untested. These limitations notwithstanding, the atlas provides a stage-resolved, spatially anchored reference against which interzone identity can be defined and in vitro systems can be scored.

## METHODS

### Timed pregnancy

All procedures performed on mice were approved by the Johns Hopkins University Animal Care and Use Committee (Protocol: MO23M290). Homozygous Sox9-IRES-GFP mice ^39^, 8 to 15 weeks old, were maintained under standard conditions (22 C, 40 to 70% humidity, 12 h light/dark cycle) with food and water ad libitum. For timed matings, one male was housed with two females for 4 h in the late afternoon and females were checked for a vaginal plug; noon of the day of the plug was taken as E0.5.

### Tissue collection for scRNA-seq

For E12.5 to E15.5 embryos and for explants, we dissected the presumptive knee by removing the autopod and the proximal femur. For E11.5 embryos, we collected the entire hindlimb. Limbs from all embryos of a litter (6 to 8) were pooled and digested in 3 mg/ml collagenase D in PBS for 1.5 h and then in 1.5 mg/ml collagenase D for 3 h. Debris was removed with a 100 um nylon filter. TER119+ and CD45+ cells were depleted with the EasySep Mouse Mesenchymal Stem/Progenitor Cell Enrichment Kit (STEMCELL Technologies) according to the manufacturer’s instructions.

### scRNA-seq library preparation and sequencing

At E11.5, E13.5, and E14.5, we loaded approximately 40,000 cells into a PIP-seq V4 T20 reaction (Illumina) to target roughly 20,000 cells per stage. At E12.5 and E15.5, we loaded approximately 200,000 cells into a PIP-seq V4 T100 reaction to target roughly 100,000 cells. For explants, we collected two replicate cultures per timepoint and loaded 20,000 cells per replicate to target roughly 10,000 cells. Libraries were sequenced to a target depth of 50,000 reads per cell. Reads were aligned with PIPseeker v3.3.0 to GRCm38 (10x Genomics Cell Ranger mouse reference 3.0.0, mm10) modified to include eGFP.

### Spatial transcriptomics

Hindlimbs were fresh frozen on liquid nitrogen in OCT and sectioned at 10um in the coronal or sagittal plane. Sections were placed on 10 mm by 10 mm Curio Seeker tiles (Curio Bioscience) for in vivo samples and 3 mm by 3 mm tiles for explants. Curio Seeker samples the whole transcriptome at 10 um bead resolution. Tile libraries were sequenced to a target depth of 2 x 10^9 reads per tile. Bead barcodes were corrected with a Python implementation of the Syrah pipeline^94^ and reads were aligned to the same eGFP-modified GRCm38 reference with the Slide-seq pipeline v3.4.2^95^. Sections were visualized with the STUF package (Cahan Lab, https://github.com/CahanLab/STUF).

### Quality control

Count matrices were processed per stage in Scanpy v1.9^48^. Mitochondrial and ribosomal fractions were computed with pySingleCellNet (pySCN)^47^. Cells were kept if they had at least 500 to 750 total UMIs and at least 1,000 detected genes, fewer than 40,000 to 65,000 total UMIs, and 1 to 15% mitochondrial reads; the exact bounds were set per dataset within these ranges based on the observed distributions. Genes detected in fewer than 0.1% of cells were removed. Ambient RNA was corrected with SoupX v1.3.0^96^ and the corrected counts were stored as a separate layer. Predicted genes, long non-coding RNAs, mitochondrial genes, ribosomal genes, and Riken genes were kept in the raw layer but excluded from downstream analyses.

SoupX-corrected counts were normalized to 10,000 counts per cell and log-transformed, and the 2,000 most variable genes were selected by the Seurat v3 method. Principal component analysis (PCA) was run on these genes. Inspection of the first 10 principal components showed separation driven by cell cycle genes and by Spidr and Bc1. When we compared samples collected at the same stage on 10x Genomics and on PIP-seq, Spidr and Bc1 were elevated in every cell type on PIP-seq, which indicates a platform artifact. We therefore computed the correlation of each principal component score with cell cycle gene set scores and with Spidr expression, using the pySCN gene set scoring functions (value-based and rank-based AUCell and UCell scores), and excluded any component whose correlation exceeded 0.15 and whose top 25 loadings included the corresponding genes.

Clustering used the pySCN parameter sweep over Leiden resolution, number of nearest neighbors, and number of principal components. We chose the parameter set that maximized the number of strict marker genes (expressed in at least 20% of cells in the cluster, at most 10% outside it, p < 0.01). Clusters were subclustered iteratively and the sweep repeated within subsets. To confirm that clusters were transcriptionally distinct, we trained a pySCN random forest classifier on 50% of cells per cluster (minimum 50 cells) and required a classification precision above 0.5; clusters below this were merged with the cluster into which their cells were most often misclassified and the test repeated. Distinct clusters were grouped into broad classes and lineages. Cell types were named from the spatial localization of their marker genes in our sections or from the literature.

### Spatial transcriptomics quality control

Per tile, we removed predicted genes, the noncoding transcript CT010467.1, and all ribosomal and mitochondrial genes, then removed spots with fewer than 15 detected genes. Within each manually defined region of interest, low-quality spots along the tissue edge were removed by applying the STUF contour profile function to total counts per spot.

| Stage | Number of sections | Total spots |
| --- | --- | --- |
| E11.5 | 7 | 89,488 |
| E12.5 | 14 | 318,996 |
| E13.5 | 12 | 340,274 |
| E14.5 | 10 | 520,120 |
| E15.5 | 4 | 270,294 |
| Explant D1 | 5 | 64,033 |
| Explant D3 | 2 | 45,998 |

### Cross dataset comparison

Published datasets^27,29,31,34,44–46^ (Figure S1A; Figure S10A) were obtained from GSE289874, GSE126966, GSE142425, GSE14441, and GSE252486 processed as above. For each stage, we trained a pySCN classifier on our annotations with a floor of 50 and a cap of 1,000 cells per type, 50% of cells for training, 50 top gene pairs, 50 top genes, and 30 principal components, on the SoupX layer, excluding types with fewer than 10 cells, and applied it to the published data of the matching stage. We also projected published cells into our PCA space and transferred labels by nearest neighbors with Scanpy’s Ingest function using the default parameterst^48,49^. A cell type was scored as present in a dataset if at least one cell was assigned. To compare annotation resolution with the integrated atlases of Zhang et al.^29^ and Herpelinck et al.^30^, we trained pySCN^47^ classifiers on the lateral plate mesoderm annotations of each atlas, restricted where possible to stylopod and zeugopod, and computed per-type precision, recall, and F1 on held-out cells.

### Stage-specific classifiers and cross-stage interzone analysis

For each of E12.5, E13.5, E14.5, and E15.5, cells were split by cell type into training and held-out sets (50% each, with a minimum of 200 and a maximum of 2,000 cells per type). From each training set we selected 2,000 highly variable genes (Seurat v3 method, batch_key = timepoint) on SoupX-corrected counts, ran PCA, and trained a pySCN classifier (nRand = 100, nTopGenes = 30, nTopGenePairs = 40, n_comps = 30). To classify published datasets (Figures S10 to S12), the classifier of the matching stage was applied and results were plotted as alluvial diagrams or bar charts.

For the cross-stage analysis, each stage classifier was applied to the held-out cells of all four stages, giving four labels per cell. Cells labeled interzone by at least one classifier were kept, and cells labeled interzone by non-consecutive stages were removed. This gave ten categories: four single-stage, three two-stage (E12.5 to E13.5, E13.5 to E14.5, E14.5 to E15.5), two three-stage (E12.5 to E14.5, E13.5 to E15.5), and one four-stage. Differentially expressed genes among the ten categories were computed at each stage (Benjamini-Hochberg adjusted p-values < 0.05), and genes were prioritized if they replicated in at least two datasets. Cell cycle-associated genes were excluded from downstream analysis to focus on lineage-specific transcriptional programs.

### Genes co-expressed with Gdf5

To identify genes co-expressed with Gdf5, we performed weighted gene co-expression network analysis using biweight midcorrelation (bicor) as the similarity metric^67^. Gene co-expression modules were identified using an automated clustering approach requiring a minimum of 5 genes per module and a minimum within-module correlation threshold of 0.4. Genes failing to meet these criteria were classified as unassigned. The module containing Gdf5 was extracted for downstream analysis to characterize the transcriptional program associated with joint interzone specification. Separately, we constructed a gene-gene similarity graph by treating genes as observations and cells as features. Analysis was restricted to highly variable genes (HVGs), and expression values were not scaled prior to similarity computation. We computed cosine similarity between genes and constructed a k-nearest neighbor graph (k = 50). Gene modules were identified by clustering this graph, while gene programs were independently extracted using non-negative matrix factorization (NMF)^74^. For each developmental stage, we recorded both the cluster module and NMF program containing Gdf5, as well as the 50 genes most similar to Gdf5 by cosine similarity. Gdf5 neighbors and Gdf5-containing modules were scored per cell using Scanpy’s score_genes function^48^ and visualized on UMAP embeddings. Statistical approaches adapted from the backend of xcell package (Cahan Lab, https://github.com/CahanLab/xcell).

### Chondrocyte differentiation pseudotime

To place in vivo and explant chondrocytes on one trajectory, we corrected batch across all samples with Harmony^97^, computed a diffusion map for chondrocytes, and selected components 2 and 3, which separated Ihh and Col10a1 expression, for trajectory inference with scFates^62^. We fit univariate smoothing splines to the fitted expression of Sox9, Sox5, Sox6, Sp7, Ihh, Panx3, and Col10a1 as functions of pseudotime and found inflection points and maxima from the first and second derivatives. Cells were assigned to four states: chondroprogenitors, before the first inflection of Sox9 (low Sox9, Sox5, Sox6; high Prrx1 and Shox2); chondrocytes, between the first inflection of Sox9 and the inflection of Sp7 (peak Sox9, Sox5, Sox6); pre-hypertrophic chondrocytes, between the Sp7 inflection points (peak Sp7, Ihh, Col10a1 rising); and hypertrophic chondrocytes, beyond the Sp7 peak, where Col10a1 reached its maximum. Pseudotime was scaled to 0 to 1 for display. Explant chondrocytes were assigned to these states with pySCN and with Ingest as above.

### Mesenchymal condensation pseudotime

To reconstruct chondrogenic differentiation trajectories, we analyzed Prrx1-expressing cells at E11.5, stratified into proximal and distal subsets. For each subset, we computed biweight midcorrelation coefficients between Sox9 and all other genes^67^, selecting the top and bottom 100 correlated genes (200 total) as a feature set enriched for chondrogenesis-associated expression variation. PCA was performed on this reduced feature space, followed by diffusion map embedding. Diffusion components capturing Sox9 expression gradients were selected for trajectory inference (components [2,3] for proximal cells; components [2, 4] for distal cells). Gene expression dynamics along pseudotime were modeled using scFates^62^, and genes were hierarchically clustered by their fitted temporal profiles using correlation distance. Gene clusters were annotated for ligand-receptor pathway membership using the CellChat database^98^.

### Axial patterning genes in spatial sections

For each section, PCA on the two-dimensional spot coordinates gave a first component approximating the proximal-distal axis and a second approximating the anterior-posterior axis. Spot positions were projected onto each component and scaled to 0 to 1. Two tests were applied per gene, adapted from the xcell package (Cahan Lab, https://github.com/CahanLab/xcell). First, expression was fit as a cubic B-spline of position with five interior knots at quantiles of the position distribution and compared to an intercept-only model by F-test. Second, positions were binned into 20 intervals and a chi-square test of independence was applied to the fraction of expressing spots per bin. P values were adjusted by the Benjamini-Hochberg procedure (FDR < 0.05) and significant genes were classed as increasing or decreasing by the sign of their correlation with position. The union of the two tests was taken per section. For in vivo stages, a gene was retained if it was significant in the same direction in at least two sections of that stage; for explants, all significant genes were retained; genes significant in opposite directions in different sections were excluded. Genes patterned at every stage were obtained by intersecting the increasing or the decreasing sets across E12.5, E13.5, E14.5, E15.5, D1, and D3.

### Reanalysis of published data

To assess the effect of Sox9 loss on Gdf5 expression, we analyzed bulk RNA-seq data from Sox9 knockout and control limb buds at E11.5 and E12.5 (GSE114522)^65^. At E11.5, Sox9 constitutive knockouts (Sox9 −/−) were compared to floxed controls (Sox9 fl/fl), while at E12.5, conditional knockouts (Sox9 fl/fl; Prx1-Cre) were compared to floxed controls. Gene expression was quantified as normalized reads per kilobase per million (NRPKM). Statistical significance was assessed using one-tailed independent samples t-tests, testing the hypothesis that Gdf5 expression is reduced in Sox9 knockouts relative to controls. P-values were reported with significance thresholds of p < 0.05 (*), p < 0.001 (**), and p < 0.0001 (***). A one-tailed test was used based on the a priori hypothesis that Sox9, as a transcriptional activator of chondrogenic genes, positively regulates Gdf5 expression; thus, Sox9 loss was predicted to reduce Gdf5 levels.

Human spatial transcriptomics sections (Figures S7, S9E, S16; Figure 4I) were obtained from Zhang et al. ^29^ and Pan et al. (GSE326326)^41^.

### In situ hybridization

Hindlimbs were fixed in 4% paraformaldehyde overnight at 4 C, washed in distilled water, incubated overnight in 30% sucrose, embedded in OCT, and stored at −80 C until cryosectioning. Hybridization used the RNAscope Multiplex Fluorescent Reagent Kit v2 (ACD) according to the manufacturer’s instructions with probes against Sox9, Gdf5, Sema3c, Igfbp5, Pamr1 on 15 um sections. Images were acquired on EVOS M7000 and Zeiss LSM 900 confocal microscope.

### Micromass culture

Whole hindlimbs from E11.5 Sox9-IRES-GFP embryos were dissociated in 0.1% trypsin and 0.1% collagenase D for 15 min at 37°C. Cells were resuspended at 100,000 cells per 5 μl in PBS and mixed with 5 μl of siRNA-Lipofectamine complex (Lipofectamine RNAiMAX, Thermo Fisher, 13778100), yielding 100,000 cells per 10 μl. Cells were seeded as 10 μl droplets in 24-well plates (n = 6 wells per condition). After 1 h at 37°C, DMEM supplemented with 10% FBS (Thermo Fisher, A4736401) and 1% antibiotic-antimycotic (Thermo Fisher, 15240062) was added gently to the side of each well. Cultures were maintained for 48 h with daily medium replacement. Knockdown of Sox9 was confirmed by RT-qPCR and flow cytometry for GFP, serving as a positive control for transfection efficiency.

After 48 h, cells were dissociated with 0.1% trypsin and 0.1% collagenase D for 5 min, and total RNA was extracted using the Monarch Total RNA Miniprep Kit (New England Biolabs). Gene expression was quantified by RT-qPCR using SYBR Green Master Mix (Bio-Rad, 1725121). Three technical replicates per gene were averaged, and relative expression was calculated using the ΔΔCt method with β-actin as the reference gene. Fold change was computed as 2^(-ΔΔCt) relative to siRNA-negative controls. Statistical significance was assessed by Welch’s t-test comparing ΔCt values between conditions, with Bonferroni correction for multiple comparisons across genes.

### Limb explant culture and siRNA injection

Hindlimbs were dissected from E12.5 embryos under sterile conditions in a biosafety cabinet, placed on permeable PET membrane supports (Falcon 353102) in 60 mm dishes (CELLTREAT 120423A001) for injection, and treated as described [58]. siRNA-Lipofectamine complexes were prepared immediately before use by mixing 1 ul of Sema3c siRNA (Thermo Fisher s73538), Igfbp5 siRNA (s68142), or Silencer Select Negative Control siRNA (4390771) with 2 ul of Lipofectamine RNAiMAX (Thermo Fisher 13778100) without dilution in Opti-MEM, pipetting gently, and incubating briefly. Approximately 100 nl of complex was loaded into a pulled glass micro-needle and injected into the knee interzone under a stereomicroscope. Explants were then placed on sterilized stainless-steel mesh (TIMESETL TXJ-277-US) in center-well organ culture dishes (Falcon 353037) containing BGJb medium (Thermo Fisher 12591038) with 1% antibiotic-antimycotic (Thermo Fisher 15240062) and cultured for 72 h with daily medium replacement and repositioning to prevent adhesion. Morphology and Sox9-GFP fluorescence were recorded on a ZEISS Stemi 508 stereomicroscope. Femoral and tibial area, length, and width were measured in ImageJ on D3 images (n =7 per condition). For each measure, groups were compared by one-way ANOVA followed by Tukey’s HSD test.

### RT-qPCR primers

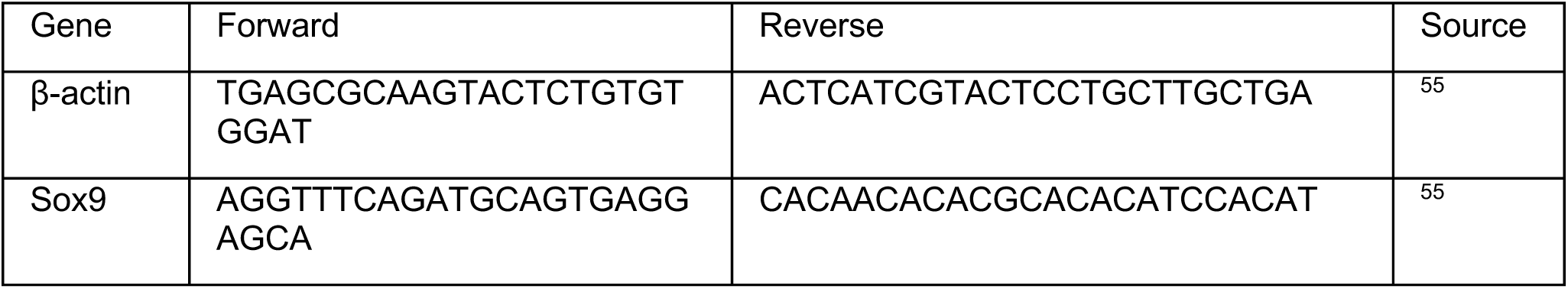
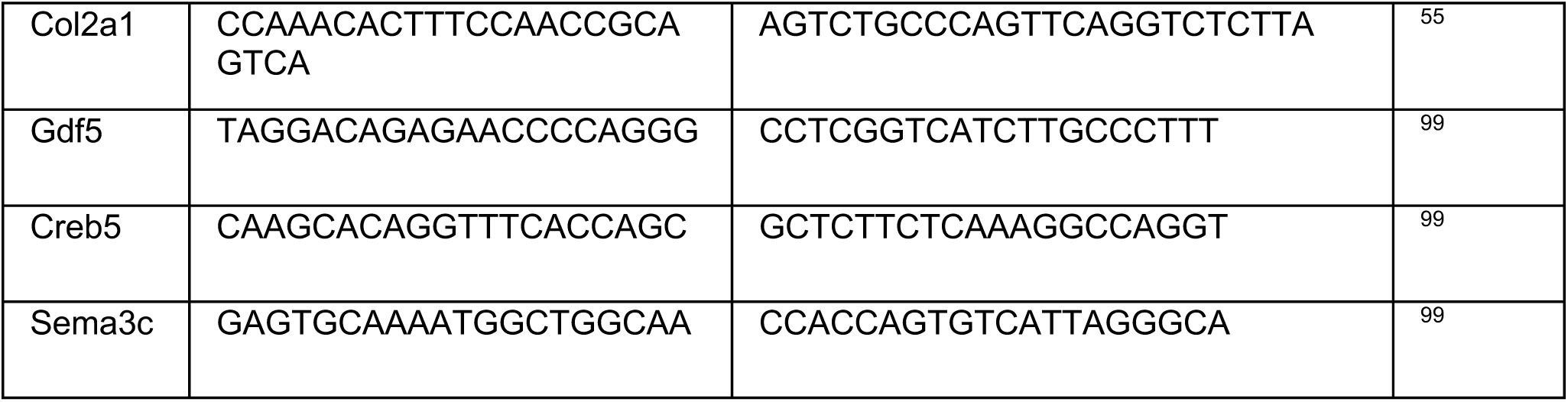

### 2siRNA

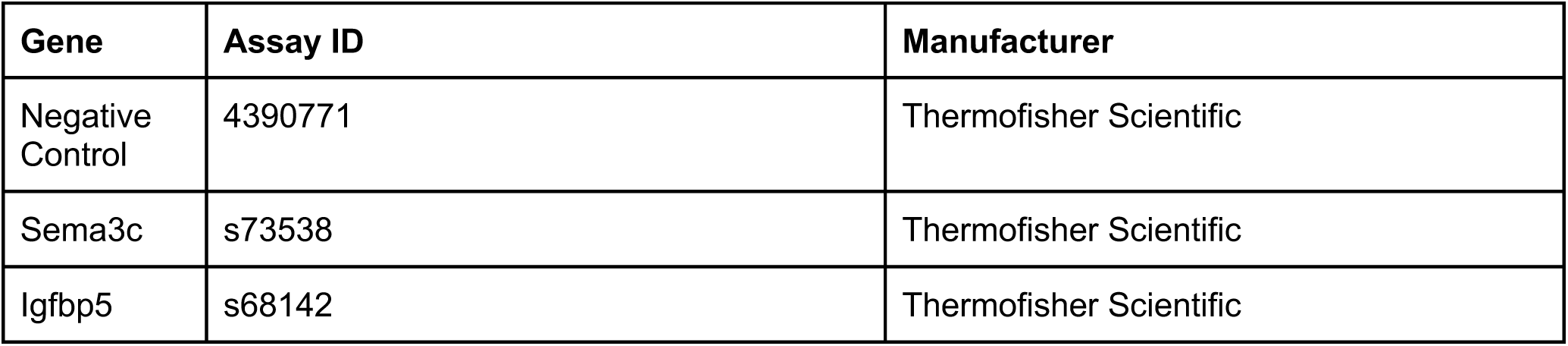

### RNAscope probes

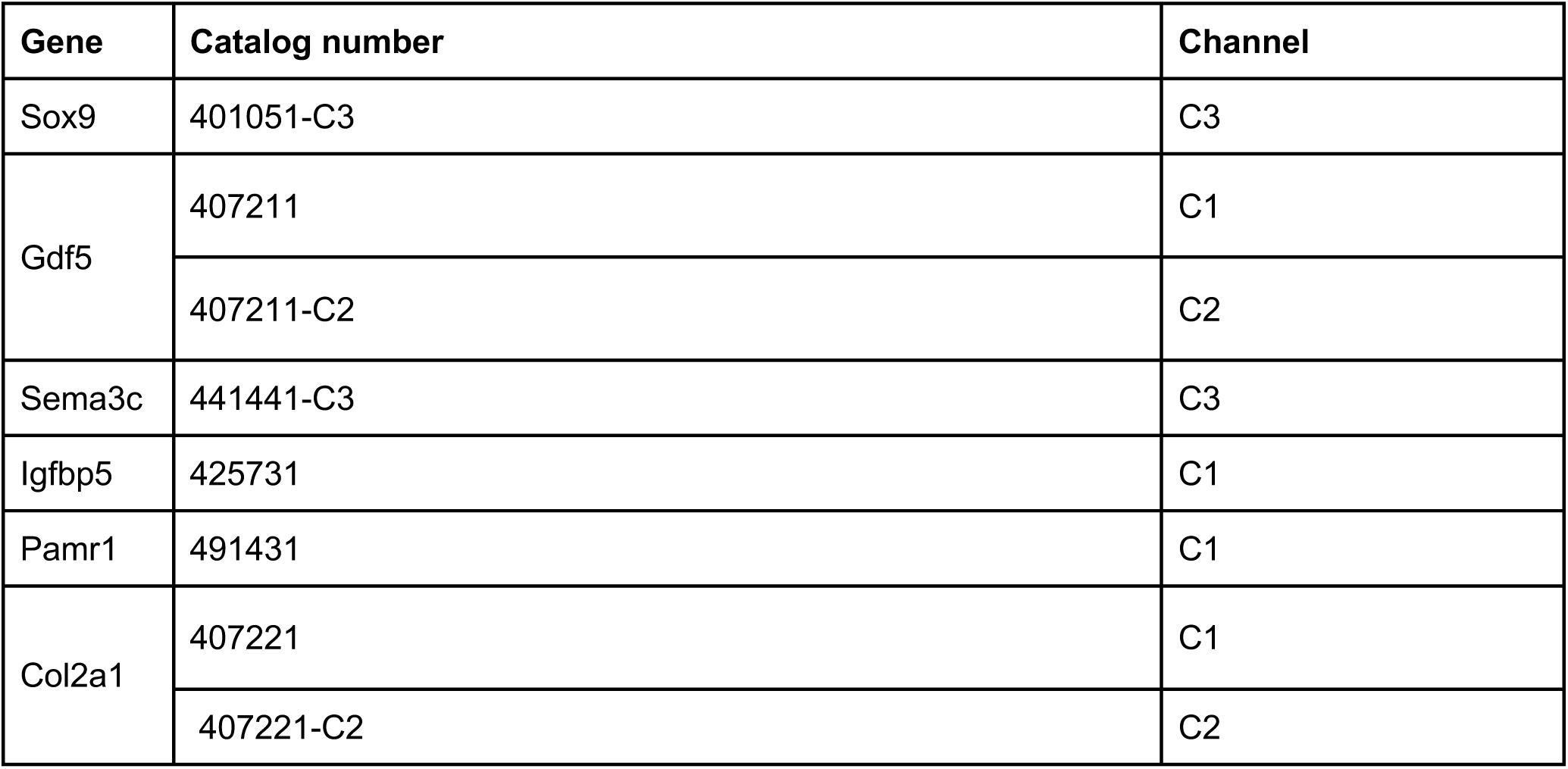

| Gene | Catalog number | Channel |
| --- | --- | --- |
| Sox9 | 401051-C3 | C3 |
| Gdf5 | 407211 | C1 |
|  | 407211-C2 | C2 |
| Sema3c | 441441-C3 | C3 |
| Igfbp5 | 425731 | C1 |
| Pamr1 | 491431 | C1 |
| Col2a1 | 407221 | C1 |
|  | 407221-C2 | C2 |

## RESOURCE AVAILABILITY

### Lead contact

Requests for further information and resources should be directed to and will be fulfilled by the lead contact, Patrick Cahan.

### Data and code availability

Single-cell RNA-seq and spatial transcriptomics data have been deposited at GEO at GSE336117 and GSE346292 and will be publicly available upon publication. Analysis scripts are available on Github (github.com/CahanLab/limb-dev-atlas), including python implementation of Syrah^94^. Microscopy data reported in this paper will be shared by the lead contact upon request. Any additional information required to reanalyze the data reported in this paper is available from the lead contact upon request.

## ACKNOWLEDGMENTS

We gratefully acknowledge Dr. Yichuan Mao, Dr. Emily Lo, Dr. Qin Bian, and Dr. Irika Sinha for their feedback and helpful discussion. We thank Dr. Aaron James for providing access to their microscope. We thank Dr. David Mohr at the Genetic Resources Core Facility, Johns Hopkins School of Medicine for scRNA-seq library sequencing and Dr. Russell Hughes and Dr. Emanuele Palescandolo at the Single Cell & Transcriptomics Core, Johns Hopkins School of Medicine, for Curio Seeker spatial transcriptomics libraries preparation and sequencing.

The authors also acknowledge funding from the National Institutes of Health (R35GM124725 to P.C.). SESO is the recipient of Predoctoral Individual National Research Service Award from National Institute of Arthritis and Musculoskeletal and Skin Diseases, National Institutes of Health (F31AR084301), and American Dissertation Fellowship from American Association of University Women (AAUW). JHY is the recipient of the Postdoctoral Research Accelerator Award from Johns Hopkins University. Some figures were created with Biorender.com and Lucid chart (lucid.co).

## AUTHORSHIP CONTRIBUTIONS

SESO, JHY, and PC conceived and designed the research studies. SESO and JHY performed the majority of experiments and acquired data, with assistance from KKY, KC, ZN, and HC. SESO, JHY, KKY, KC, and HC performed colony maintenance, timed pregnancies, embryo isolation, cryo-sectioning, RNAscope, and confocal microscopy. SESO performed micromass culture experiments. SESO and ZN performed RT-qPCR for micromass culture experiments. JHY performed ex-vivo explant cultures and semi-quantitative grading for knockdown phenotype. SESO performed single cell library preparations. JHY performed fresh-frozen sectioning for spatial transcriptomics. SESO performed scRNA-seq and statistical analysis with guidance from PC, JHY, ZN, and HC. SESO wrote the manuscript with support from JHY and PC. PC supervised the study. All authors reviewed and approved the final manuscript.

## DECLARATION OF GENERATIVE AI AND AI-ASSISTED TECHNOLOGIES

During the preparation of this work, the authors used Claude-opus-4.5 to troubleshoot code, including analyzing error messages and checking syntax errors, and flag conflicting information between the manuscript and figure legends. Claude Opus 5.5 and Fable 5.1 (Anthropic) were used when writing the manuscript to improve conciseness and clarity. After using these tools, the authors reviewed and edited the content as needed and take full responsibility for the manuscript.

## Supplementary figure legends

**Figure S1. Comparison of the atlas to published scRNA-seq datasets of the developing limb.** Related to Figure 1. (A) Datasets used for comparison, by stage. Blue, cells; pink, spots. Brackets mark the datasets to which the Zhang et al. (orange) and Herpelinck et al. (green) integrated annotations were applied.

(B) Presence of each cell type defined in this study (rows) in each published dataset (columns) at the same stage, as determined by pySCN and by Ingest classification. Green, detected; white, not detected. The fraction of cells of each type in this study is given at right.

(C) Precision, recall, and F1 score of pySCN classifiers trained on the lateral plate mesoderm annotations of this study (Onggo), of Zhang et al., and of Herpelinck et al., evaluated on held-out cells at each stage. Each point is one cell type; the number of cell types is given above each box. Neither integrated atlas has an annotated E14.5 stage.

**Figure S2. Annotation of the E11.5 hindlimb.** Related to Figure 1. (A) Sox9-GFP in a whole E11.5 hindlimb. Scale bar, 200 um.

(B) Dot plot of marker genes (columns) for each cell type (rows), grouped by broad class. Dot size, fraction of cells expressing; color, mean scaled expression.

(C) Alluvial diagram relating lineage, broad class, and cell type.

(D) Spatial expression of regional and lineage markers (red) over the pre-cartilage condensation (Sox9, Wwp2, Acan; grey): Isl1; Pdgfra, Col1a1, Col3a1; Gdf5; Meis1, Meis2; Hoxa11; Hoxd12, Hoxd13; Hand2; Zic3, Zic5; Fgf8.

**Figure S3. Annotation of the E12.5 hindlimb.** Related to Figure 1. (A) Sox9-GFP in a whole E12.5 hindlimb. Scale bar, 200 um.

(B) Dot plot of marker genes for each cell type, as in Figure S2B.

(C) Alluvial diagram relating lineage, broad class, and cell type.

(D) Spatial expression of regional, interzone, and lineage markers (red) over the cartilage template (Sox9, Wwp2, Acan; grey): Isl1; Hoxa11; Ihh, Panx3; Hoxc11; Apcdd1; Pamr1; Pdgfra, Dlk1, Cxcl12, Dkk2; Tbx18, Shox2, Cadm2, Foxp2, Gas2, Col8a1; Gdf5, Osr2, Sulf1, Sfrp2; Scx, Mkx, Tnmd; Hoxa11, Sall1, Zic3, Meox2; Pamr1, Tnc, Ncam1, Cdh2.

(E) Spatial expression of Tbx5, Hoxc11, Sall1, and Zic3 (red) over the cartilage template (grey).

**Figure S4. Annotation of the E13.5 hindlimb.** Related to Figure 1. (A) Sox9-GFP in a whole E13.5 hindlimb. Scale bar, 200 um.

(B) Dot plot of marker genes for each cell type, as in Figure S2B.

(C) Alluvial diagram relating lineage, broad class, and cell type.

(D) Spatial expression of markers (red) over the cartilage template (Sox9, Wwp2, Acan; grey): Pdgfra, Dlk1, Cxcl12, Dkk2; Apcdd1, Axin2, Twist2; Alpl, Dlx5, Sp7; Gdf5, Osr2, Sulf1, Sfrp2, Plxna4; Barx1, Scrg1, Tbx5; Ihh, Panx3; Mkx, Tnmd; Pitx2, Cytl1, Fxyd7, Alx1, Susd5, Kcne4; Htra1, Adgrg2; Zic3, Scara5, Bdb2, Chsy3, Spry2, Hoxd9.

(E) Spatial expression of tendon and ligament genes Htra1, Adgrg2, Prss35, Kera, and Chodl (red) over the cartilage template (grey).

(F) Spatial expression of Hoxc11, Sall1, Zic3, and Tbx5 (red) over the cartilage template (grey).

**Figure S5. Annotation of the E14.5 hindlimb.** Related to Figure 1. (A) Sox9-GFP in a whole E14.5 hindlimb. Scale bar, 200 um.

(B) Dot plot of marker genes for each cell type, as in Figure S2B.

(C) Alluvial diagram relating lineage, broad class, and cell type.

(D) Spatial expression of markers (red) over the cartilage template (Sox9, Wwp2, Acan; grey): Pdgfra, Dlk1, Cxcl12, Dkk2; Ihh, Panx3; Col10a1; Gdf5, Osr2, Sulf1, Sfrp2, Plxna4; Mkx, Tnmd; Alpl, Dlx5, Sp7; Apcdd1, Col23a1, Axin2, Twist2, Apela; Apod, Dpt, Ren1, Lgals9, Osr1, Abi3bp; Kera.

**Figure S6. Annotation of the E15.5 hindlimb.** Related to Figure 1. (A) Sox9-GFP in a whole E15.5 hindlimb. Scale bar, 200 um.

(B) Dot plot of marker genes for each cell type, as in Figure S2B.

(C) Alluvial diagram relating lineage, broad class, and cell type.

(D) Spatial expression of markers (red) over the cartilage template (Sox9, Wwp2, Acan; grey): Pdgfra, Dlk1, Cxcl12, Dkk2; Apod, Dpt, Ren1, Osr1, Abi3bp; Ihh, Panx3; Col10a1; Spp1, Tnfrsf11b, Bglap; Gdf5; Aspn, Eln, Col8a1; Alpl, Dlx5, Sp6, Ibsp.

(E) Tendon and ligament genes at E15.5. Top row, gene sets: Mkx, Tnmd; Htra1, Aqp1, Ssc5d, Dkk3, Sfrp2; Kera, Col14a1, Cpm. Below, single genes: Chodl, Clip, Htra3, Zfp185, Thbs4, Prss35, Kera, Col14a1, Cpm, Htra1, Aqp1, Ssc5d, Dkk3, Sfrp2. Red, the indicated genes; grey, cartilage template.

**Figure S7. Tendon and ligament markers in the human hindlimb at 8 PCW.** Related to Figure 1. MKX, TNMD, PRSS35, and SSC5D (blue) over the cartilage template (COL9A1, ACAN, WWP2; red) in published spatial transcriptomics sections from Zhang et al. (two sections, left) and Pan et al. (two sections, right).

**Figure S8. Annotation of explant cultures and axial patterning genes.** Related to Figure 2. (A) D1 explant. Left, alluvial diagram relating lineage, broad class, and cell type. Right, dot plot of marker genes for each cell type. (B) D3 explant, as in (A). (C) Expression (left of each pair) and detection rate (right) of Meis1, Meis2, Shox2, Hoxa13, Hoxd10, Hoxd11, Hoxd12, Hoxd13, and Msx1 as a function of position along PC1 (proximal to distal, scaled 0 to 1) in each spatial section, colored by stage (E12.5, E13.5, E14.5, E15.5, D1, D3). Shaded bands, 95% confidence interval of the fit. (D) As in (C) for Hoxc11, Hand2, Kera, and Tbx5 along PC2 (anterior to posterior, scaled 0 to 1).

**Figure S9. Mesenchymal condensation trajectories at E11.5 and Sema3c in the human knee.** Related to Figure 3. (A) Gdf5 expression (normalized RPKM) in E12.5 limbs from Sox9 fl/fl and Sox9 fl/fl; Prrx1-Cre embryos, from Liu et. al published RNA-seq data. n = 3 per genotype. *, p < 0.05.

(B) UMAP of E11.5 Prrx1+ lateral plate mesoderm colored by Sox9, Col2a1, Col1a1, Sema3c, and Gdf5 expression, and by trajectory group (proximal, distal, or not assigned). Right, dot plot of Sox9, Sox5, Sox6, Acan, Sema3c, Gdf5, Col1a1, Col2a1, Meis1, Meis2, Hoxd12, and Hoxd13 in the proximal and distal groups.

(C) Fitted expression along scaled pseudotime of genes that change along the proximal trajectory, grouped by pathway or protein family (left color bar). Genes referred to in the text are in red.

(D) As in (C) for the distal trajectory.

(E) Co-expression analysis of Sox9, Gdf5, and Sema3c at E11.5. Top Left: Distribution of the number of genes (Sox9, Gdf5, Sema3c) expressed per cell. Percentages indicate the proportion of total cells in each category. Top Middle: Percentage of cells co-expressing all three genes (Sox9, Gdf5, and Sema3c) by cell type. Top Right: Heatmap showing the percentage of cells expressing each gene within each cell type.

(F) Left: UMAP embedding colored by the number of genes expressed per cell (0–3). Middle: UMAP embedding highlighting cells co-expressing all three genes (red) against other cells (gray). Right: UMAP embedding showing cell type annotations for reference. Co-expression was defined as expression above zero UMIs.

(G) SEMA3C (blue) over the cartilage template (ACAN, COL9A1, WWP2; red) in published spatial transcriptomics sections of the human hindlimb at 6.5, 7.1, and 8.1 PCW from Zhang et. al.

**Figure S10. Fraction of interzone cells in published datasets with and without Gdf5-lineage enrichment.** Related to Figure 4. (A) Published datasets classified, with stage, strain, and enrichment.

(B) Color key for cell types in (C).

(C) Left, composition of the limb mesenchyme of each dataset at E12.5, E13.5, E14.5, and E15.5 as classified by the stage-specific pySCN classifier trained on this study. Right, the percentage of cells classified as interzone (IZ, IZ chondrogenic, IZ talocrural, or IZ intra-articular, where present) in each dataset.

**Figure S11. Classification of E11.5 chondroprogenitors by the E12.5 classifier.** Related to Figure 4. Alluvial diagrams relating the E11.5 cell type (left) to the label assigned by the E12.5 pySCN classifier (right) for E11.5 cells from this study (Sox9-IRES-GFP), from Kelly et al. (C57BL/6J), and from Matsushita et al. (Prrx1-Cre lineage).

**Figure S12. Cross-stage classification of interzone cells.** Related to Figure 4. (A) Alluvial diagrams for cells of each stage (E12.5, E13.5, E14.5, E15.5; rows), relating the labels assigned by the E12.5, E13.5, E14.5, and E15.5 classifiers (columns, left to right).

(B) UMAP of the E13.5 Col2a1-lineage dataset of Matsushita et al. colored by the authors’ clusters and by Sox9, Col2a1, and Gdf5 expression.

(C) Dot plot of Sox9, Gdf5, Barx1, Col2a1, and Scrg1 in each author cluster. Cluster 4 is Gdf5-high and Col2a1-high.

(D) Alluvial diagram relating author clusters 4 and 9 to the labels assigned by the E12.5 and E13.5 classifiers.

**Figure S13. Genes co-expressed with Gdf5 across stages.** Related to Figure 4. (A) Biweight midcorrelation with Gdf5 at E12.5, E13.5, E14.5, and E15.5 for every gene whose correlation exceeded threshold at any stage. Grey, below threshold. Figure 4D shows the subset referred to in the text.

(B) At E12.5 and E13.5: Col2a1 expression; the correlation module containing Gdf5; the NMF program containing Gdf5; and the 50 genes most similar to Gdf5 by correlation and by k-nearest neighbors, scored per cell on the UMAP (as in Figure 4E).

**Figure S14. Core interzone markers in spatial sections at E12.5 and E13.5.** Related to Figure 4. Gdf5, Cntn4, Igfbp5, Ism1, Man1a, Pamr1, Pdlim3, Plxna4, Prkd1, Prss23, Sema3c, and Tafa2 (red) over the cartilage template (Col9a1, Acan, Wwp2; grey) in three sections at E12.5 and three at E13.5.

**Figure S15. Core interzone markers in spatial sections at E14.5 and E15.5.** Related to Figure 4. As in Figure S14 for three sections at E14.5 and three at E15.5.

**Figure S16. Human homologs of the core interzone markers in the developing hindlimb.** Related to Figure 4. Expression of the human homologs (red) over the cartilage template (COL9A1, ACAN, WWP2; blue) in published spatial transcriptomics sections at PCW 6 (Zhang et al., three sections; Pan et al., two sections) and PCW 7 (Zhang et al., four sections).

