## Supplemental Figures for "A spatial and single-cell transcriptomic atlas of the embryonic synovial joint and its explant model"

### Supp. 1

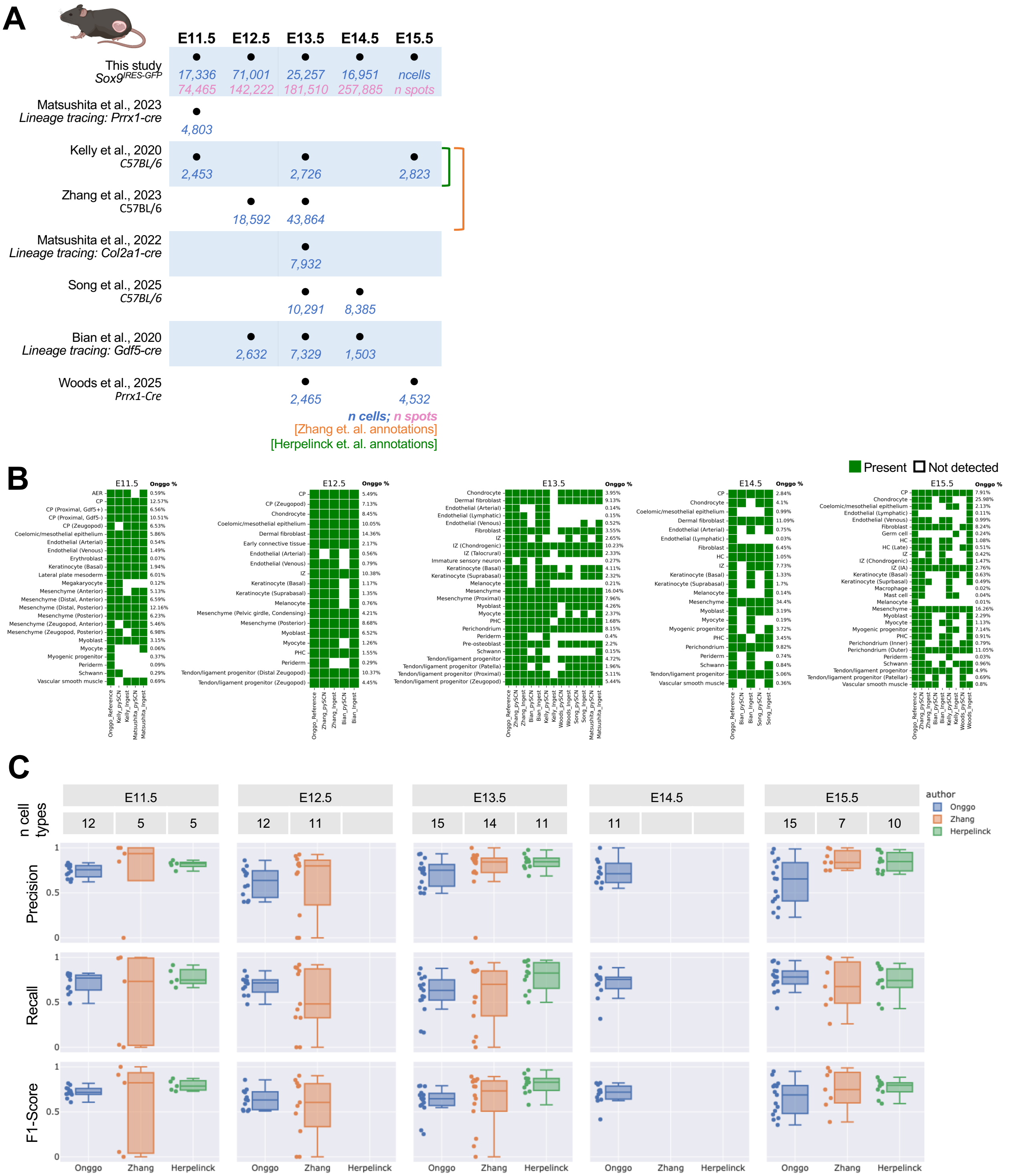
A
E11.5
E12.5
E13.5
E14.5
E15.5
This study
Sox9IRES-GFP
17,336
71,001
25,257
16,951
ncells
74,465
142,222
181,510
257,885
n spots
Matsushita et al., 2023
Lineage tracing: Prrx1-cre
4,803
Kelly et al., 2020
C57BL/6
2,453
2,726
2,823
Zhang et al., 2023
C57BL/6
18,592
43,864
Matsushita et al., 2022
Lineage tracing: Col2a1-cre
7,932
Song et al., 2025
C57BL/6
10,291
8,385
Bian et al., 2020
Lineage tracing: Gdf5-cre
2,632
7,329
1,503
Woods et al., 2025
Prrx1-Cre
2,465
4,532
n cells; n spots
[Zhang et. al. annotations]
[Herpelinck et. al. annotations]
B
Present
Not detected
E11.5
E12.5
E13.5
E14.5
E15.5
Onggo %
C
E11.5
E12.5
E13.5
E14.5
E15.5
n cell types
author
Onggo
Zhang
Herpelinck
Precision
Recall
F1-Score
Onggo
Zhang
Herpelinck

### Supp. 2

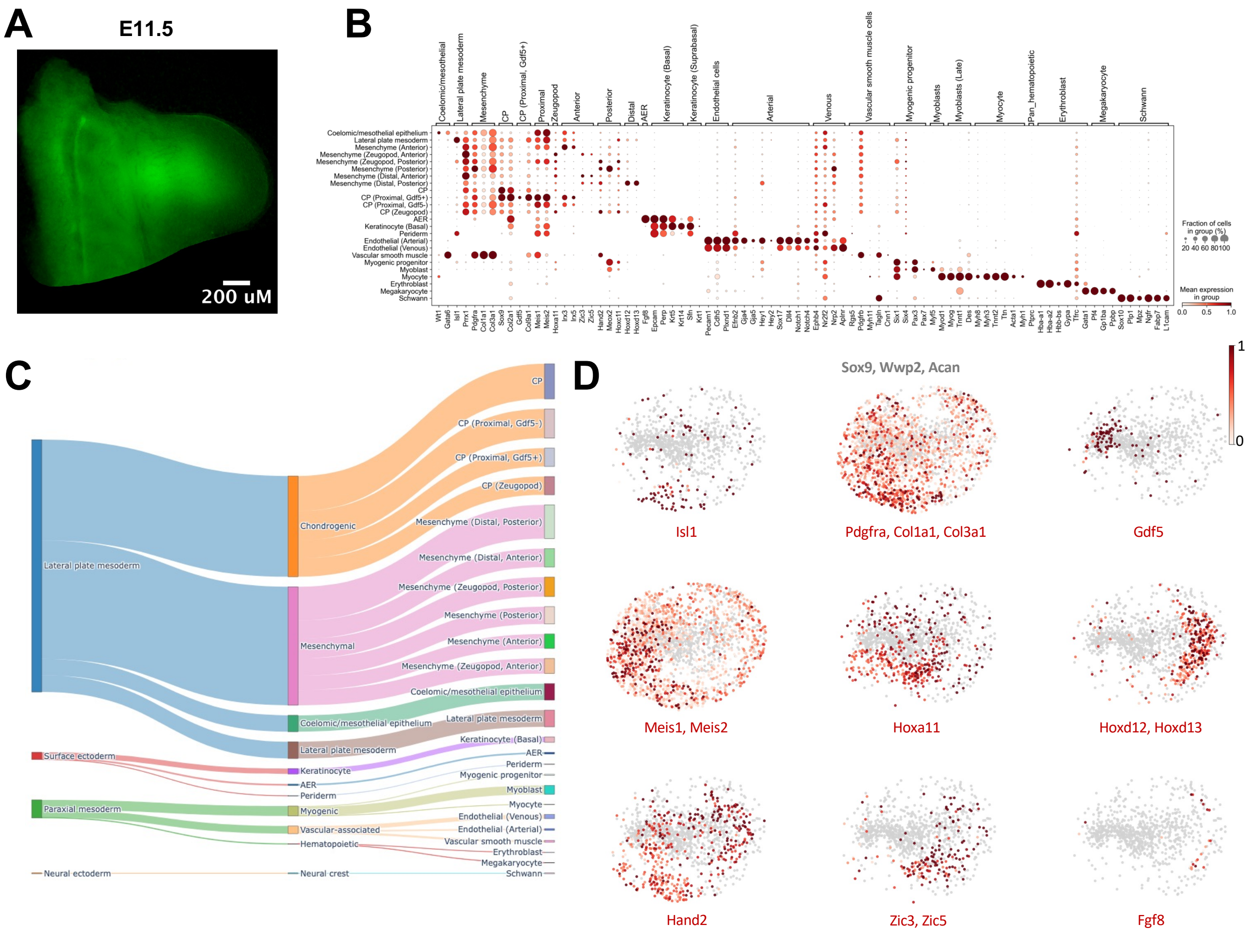
A
E11.5
200 uM
B
Fraction of cells in group (%)
20 40 60 80 100
Mean expression in group
0.0 0.5 1.0
C
Lateral plate mesoderm
Surface ectoderm
Paraxial mesoderm
Neural ectoderm
Chondrogenic
Mesenchymal
Coelomic/mesothelial epithelium
Lateral plate mesoderm
Keratinocyte
AER
Periderm
Myogenic
Vascular-associated
Hematopoietic
Neural crest
CP
CP (Proximal, Gdf5-)
CP (Proximal, Gdf5+)
CP (Zeugopod)
Mesenchyme (Distal, Posterior)
Mesenchyme (Distal, Anterior)
Mesenchyme (Zeugopod, Posterior)
Mesenchyme (Posterior)
Mesenchyme (Anterior)
Mesenchyme (Zeugopod, Anterior)
Coelomic/mesothelial epithelium
Lateral plate mesoderm
Keratinocyte (Basal)
AER
Periderm
Myogenic progenitor
Myoblast
Myocyte
Endothelial (Venous)
Endothelial (Arterial)
Vascular smooth muscle
Erythroblast
Megakaryocyte
Schwann
D
Sox9, Wwp2, Acan
1
0
Isl1
Pdgfra, Col1a1, Col3a1
Gdf5
Meis1, Meis2
Hoxa11
Hoxd12, Hoxd13
Hand2
Zic3, Zic5
Fgf8

Supp. 3

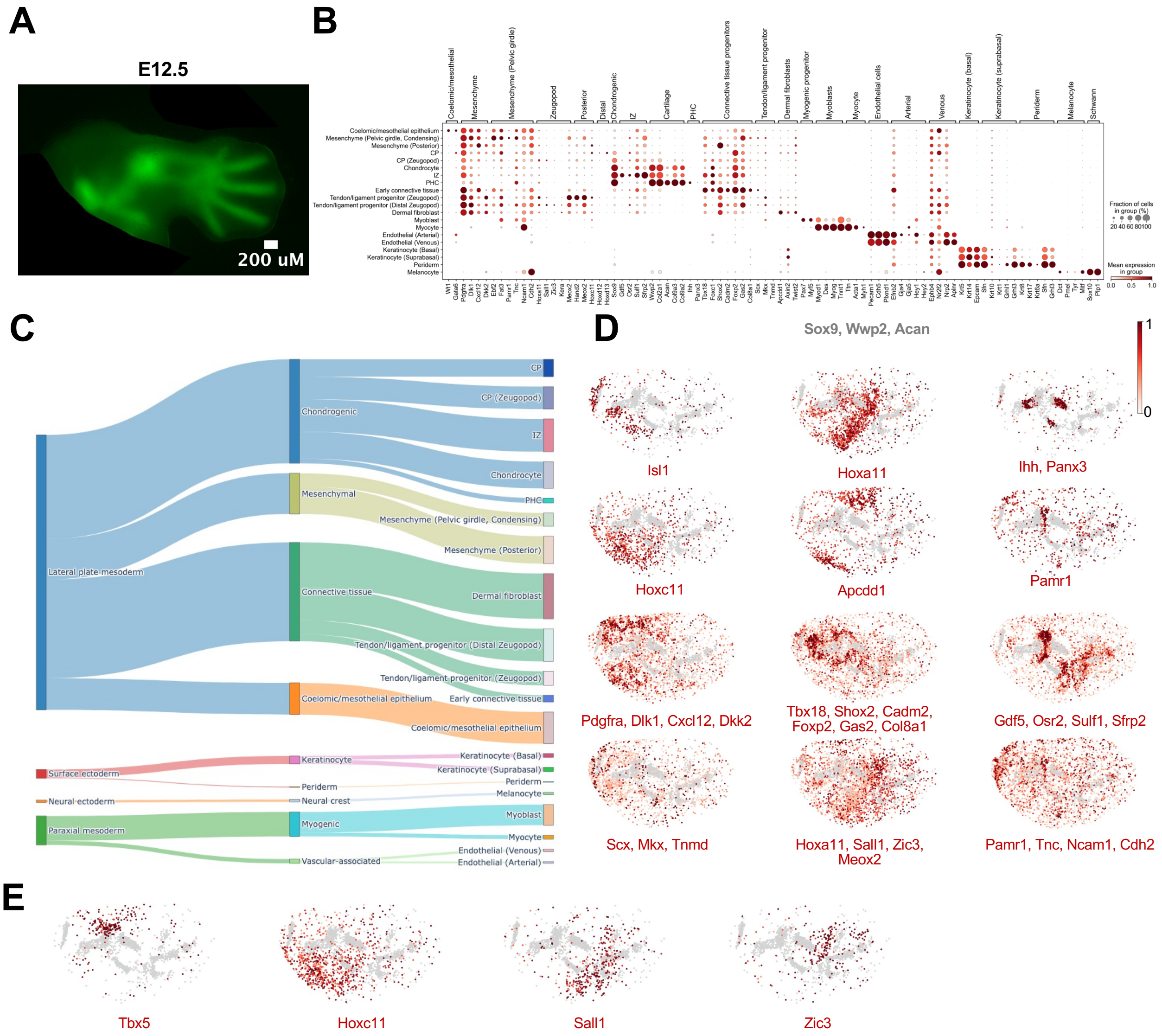

A
E12.5
200 uM
B
Coelomic/mesothelial epithelium
Mesenchyme (Pelvic girdle, Condensing)
Mesenchyme (Posterior)
CP
CP (Zeugopod)
Chondrocyte
IZ
PHC
Early connective tissue
Tendon/ligament progenitor (Zeugopod)
Tendon/ligament progenitor (Distal Zeugopod)
Dermal fibroblast
Myoblast
Myocyte
Endothelial (Arterial)
Endothelial (Venous)
Keratinocyte (Basal)
Keratinocyte (Suprabasal)
Periderm
Melanocyte
Fraction of cells in group (%)
20 40 60 80 100
Mean expression in group
0.0 0.5 1.0
C
Lateral plate mesoderm
Surface ectoderm
Neural ectoderm
Paraxial mesoderm
Chondrogenic
Mesenchymal
Connective tissue
Coelomic/mesothelial epithelium
Keratinocyte
Periderm
Neural crest
Myogenic
Vascular-associated
CP
CP (Zeugopod)
IZ
Chondrocyte
PHC
Mesenchyme (Pelvic girdle, Condensing)
Mesenchyme (Posterior)
Dermal fibroblast
Tendon/ligament progenitor (Distal Zeugopod)
Tendon/ligament progenitor (Zeugopod)
Early connective tissue
Coelomic/mesothelial epithelium
Keratinocyte (Basal)
Keratinocyte (Suprabasal)
Periderm
Melanocyte
Myoblast
Myocyte
Endothelial (Venous)
Endothelial (Arterial)
D
Sox9, Wwp2, Acan
1
0
Isl1
Hoxa11
Ihh, Panx3
Hoxc11
Apcdd1
Pamr1
Pdgfra, Dlk1, Cxcl12, Dkk2
Tbx18, Shox2, Cadm2, Foxp2, Gas2, Col8a1
Gdf5, Osr2, Sulf1, Sfrp2
Scx, Mkx, Tnmd
Hoxa11, Sall1, Zic3, Meox2
Pamr1, Tnc, Ncam1, Cdh2
E
Tbx5
Hoxc11
Sall1
Zic3

**Supp. 4**

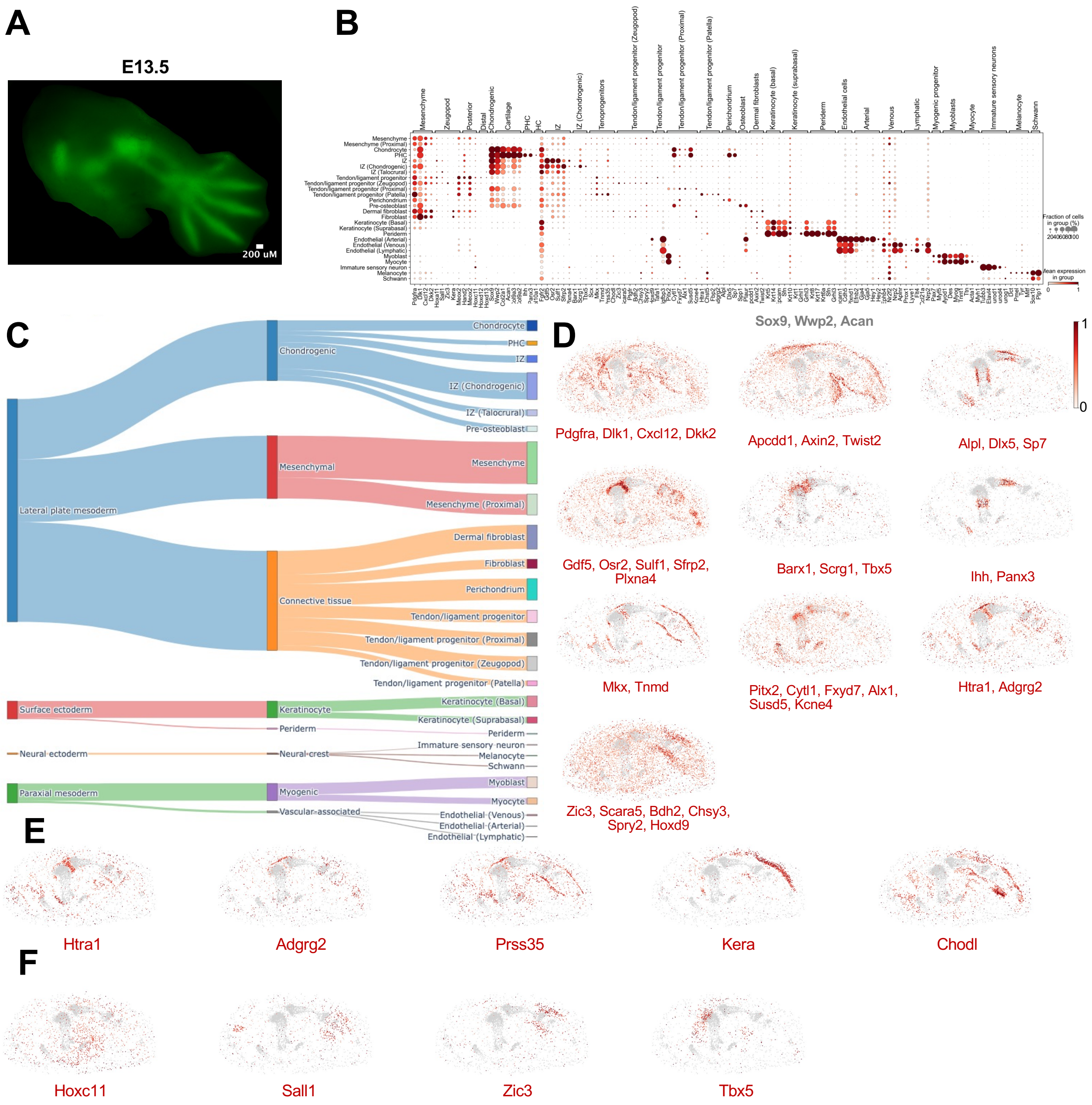

A
E13.5
200 uM
B
C
Lateral plate mesoderm
Chondrogenic
Chondrocyte
PHC
IZ
IZ (Chondrogenic)
IZ (Talocrural)
Pre-osteoblast
Mesenchymal
Mesenchyme
Mesenchyme (Proximal)
Connective tissue
Dermal fibroblast
Fibroblast
Perichondrium
Tendon/ligament progenitor
Tendon/ligament progenitor (Proximal)
Tendon/ligament progenitor (Zeugopod)
Tendon/ligament progenitor (Patella)
Surface ectoderm
Keratinocyte
Keratinocyte (Basal)
Keratinocyte (Suprabasal)
Periderm
Neural ectoderm
Neural crest
Immature sensory neuron
Melanocyte
Schwann
Paraxial mesoderm
Myogenic
Myoblast
Myocyte
Vascular-associated
Endothelial (Venous)
Endothelial (Arterial)
Endothelial (Lymphatic)
D
Sox9, Wwp2, Acan
1
0
Pdgfra, Dlk1, Cxcl12, Dkk2
Apcdd1, Axin2, Twist2
Alpl, Dlx5, Sp7
Gdf5, Osr2, Sulf1, Sfrp2, Plxna4
Barx1, Scrg1, Tbx5
Ihh, Panx3
Mkx, Tnmd
Pitx2, Cytl1, Fxyd7, Alx1, Susd5, Kcne4
Htra1, Adgrg2
Zic3, Scara5, Bdh2, Chsy3, Spry2, Hoxd9
E
Htra1
Adgrg2
Prss35
Kera
Chodl
F
Hoxc11
Sall1
Zic3
Tbx5

### Supp. 5

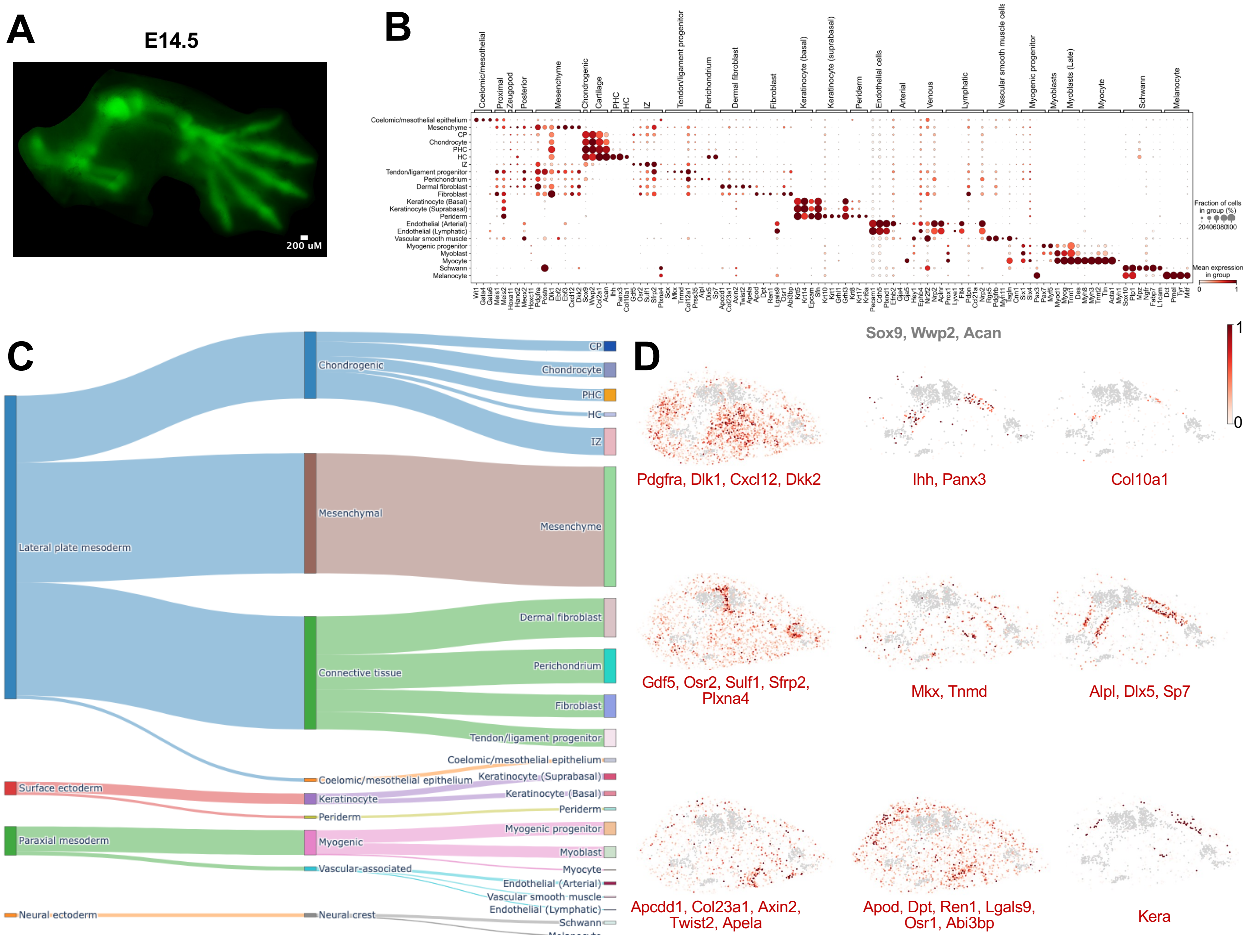

A
E14.5
200 uM
B
C
Lateral plate mesoderm
Surface ectoderm
Paraxial mesoderm
Neural ectoderm
Chondrogenic
Mesenchymal
Connective tissue
Coelomic/mesothelial epithelium
Keratinocyte
Periderm
Myogenic
Vascular-associated
Neural crest
CP
Chondrocyte
PHC
HC
IZ
Mesenchyme
Dermal fibroblast
Perichondrium
Fibroblast
Tendon/ligament progenitor
Coelomic/mesothelial epithelium
Keratinocyte (Suprabasal)
Keratinocyte (Basal)
Periderm
Myogenic progenitor
Myoblast
Myocyte
Endothelial (Arterial)
Vascular smooth muscle
Endothelial (Lymphatic)
Schwann
D
Sox9, Wwp2, Acan
1
0
Pdgfra, Dlk1, Cxcl12, Dkk2
Ihh, Panx3
Col10a1
Gdf5, Osr2, Sulf1, Sfrp2, Plxna4
Mkx, Tnmd
Alpl, Dlx5, Sp7
Apcdd1, Col23a1, Axin2, Twist2, Apela
Apod, Dpt, Ren1, Lgals9, Osr1, Abi3bp
Kera

### Supp. 6

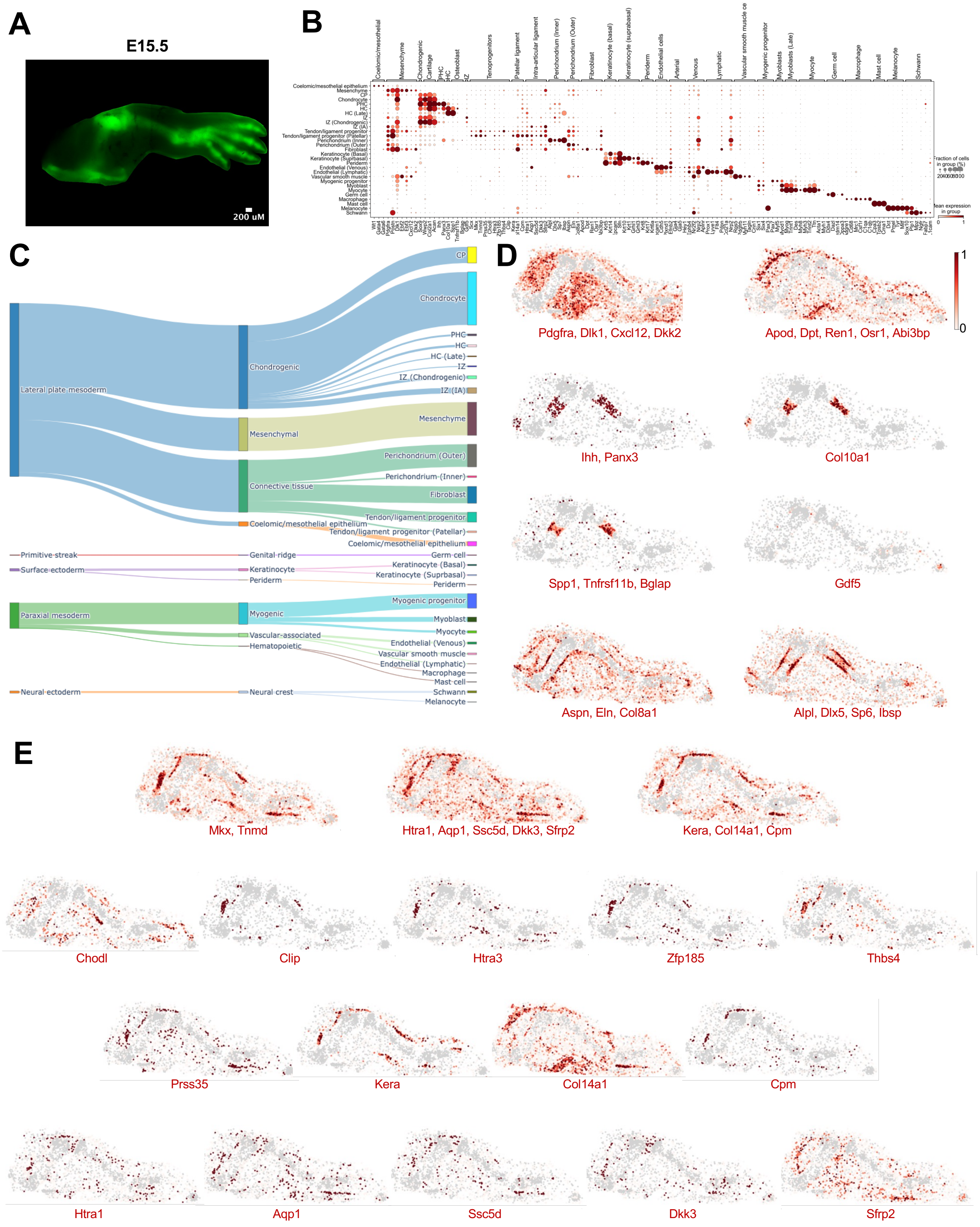
A
E15.5
200 uM
B
C
Lateral plate mesoderm
Chondrogenic
Mesenchymal
Connective tissue
Coelomic/mesothelial epithelium
CP
Chondrocyte
PHC
HC
HC (Late)
IZ
IZ (Chondrogenic)
IZ (IA)
Mesenchyme
Perichondrium (Outer)
Perichondrium (Inner)
Fibroblast
Tendon/ligament progenitor
Tendon/ligament progenitor (Patellar)
Coelomic/mesothelial epithelium
Primitive streak
Genital ridge
Germ cell
Surface ectoderm
Keratinocyte
Keratinocyte (Basal)
Keratinocyte (Suprbasal)
Periderm
Periderm
Paraxial mesoderm
Myogenic
Myogenic progenitor
Myoblast
Myocyte
Vascular-associated
Hematopoietic
Endothelial (Venous)
Vascular smooth muscle
Endothelial (Lymphatic)
Macrophage
Mast cell
Neural ectoderm
Neural crest
Schwann
Melanocyte
D
1
0
Pdgfra, Dlk1, Cxcl12, Dkk2
Apod, Dpt, Ren1, Osr1, Abi3bp
Ihh, Panx3
Col10a1
Spp1, Tnfrsf11b, Bglap
Gdf5
Aspn, Eln, Col8a1
Alpl, Dlx5, Sp6, Ibsp
E
Mkx, Tnmd
Htra1, Aqp1, Ssc5d, Dkk3, Sfrp2
Kera, Col14a1, Cpm
Chodl
Clip
Htra3
Zfp185
Thbs4
Prss35
Kera
Col14a1
Cpm
Htra1
Aqp1
Ssc5d
Dkk3
Sfrp2

#### Supp. 7

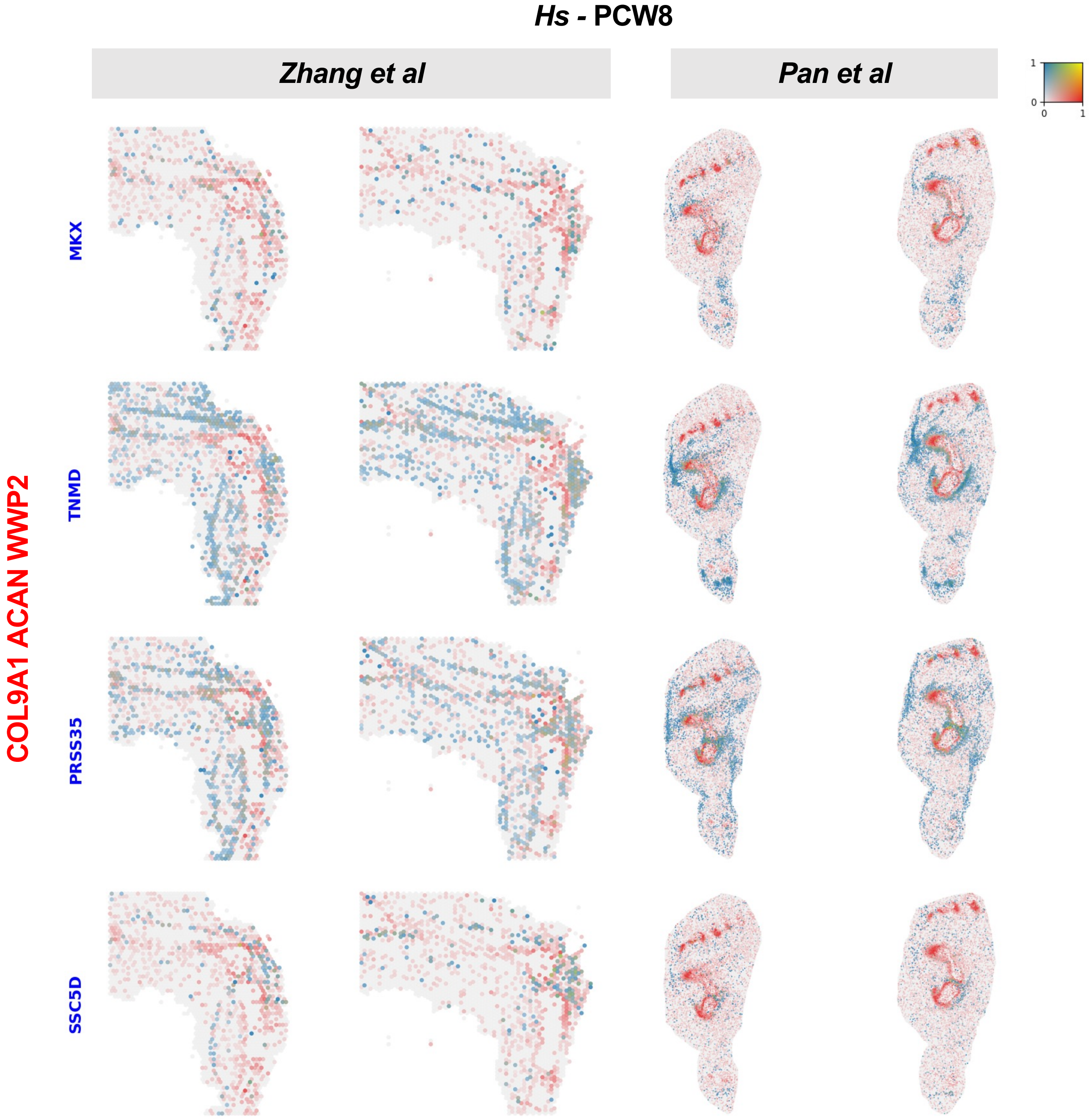

*Hs* - PCW8
*Zhang et al*
*Pan et al*
COL9A1 ACAN WWP2
MKX
TNMD
PRSS35
SSC5D
1
0
0
1

### Supp. 8

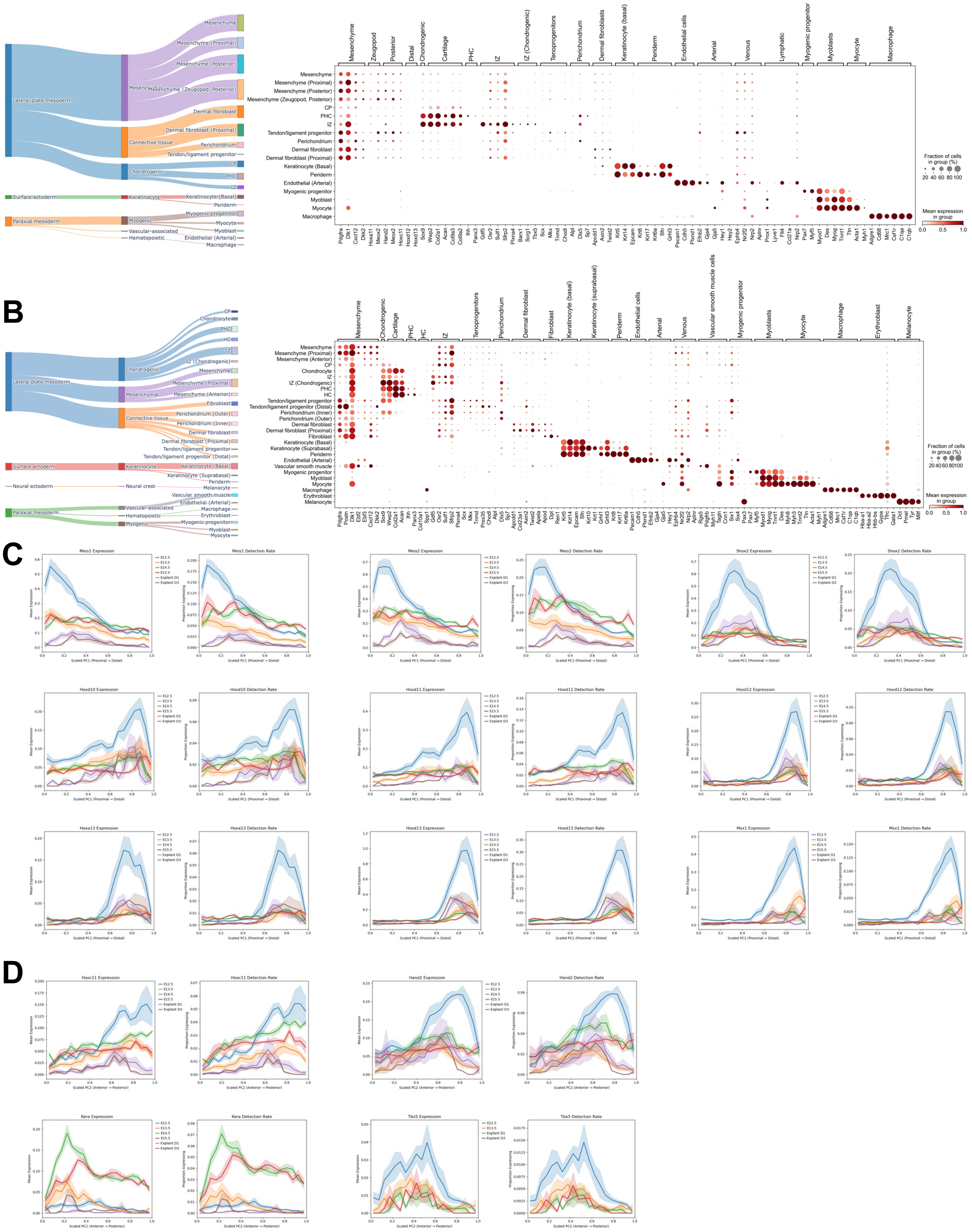
A
Lateral plate mesoderm
Surface ectoderm
Paraxial mesoderm
Connective tissue
Chondrogenic
Keratinocyte
Myogenic
Vascular-associated
Hematopoietic
Mesenchyme
Mesenchyme (Proximal)
Mesenchyme (Posterior)
Mesenchyme (Zeugopod, Posterior)
Dermal fibroblast
Dermal fibroblast (Proximal)
Perichondrium
Tendon/ligament progenitor
CP
PHC
IZ
Keratinocyte (Basal)
Periderm
Myogenic progenitor
Myocyte
Myoblast
Endothelial (Arterial)
Macrophage
Fraction of cells in group (%)
Mean expression in group
B
Neural ectoderm
Neural crest
Chondrocyte
HC
IZ (Chondrogenic)
Mesenchyme (Anterior)
Fibroblast
Perichondrium (Outer)
Perichondrium (Inner)
Tendon/ligament progenitor (Distal)
Keratinocyte (Suprabasal)
Melanocyte
Vascular smooth muscle
Erythroblast
C
Meis1 Expression
Meis1 Detection Rate
Meis2 Expression
Meis2 Detection Rate
Shox2 Expression
Shox2 Detection Rate
Hoxd10 Expression
Hoxd10 Detection Rate
Hoxd11 Expression
Hoxd11 Detection Rate
Hoxd12 Expression
Hoxd12 Detection Rate
Hoxa13 Expression
Hoxa13 Detection Rate
Hoxd13 Expression
Hoxd13 Detection Rate
Msx1 Expression
Msx1 Detection Rate
Scaled PC1 (Proximal → Distal)
D
Hoxc11 Expression
Hoxc11 Detection Rate
Hand2 Expression
Hand2 Detection Rate
Kera Expression
Kera Detection Rate
Tbx5 Expression
Tbx5 Detection Rate
Scaled PC2 (Anterior → Posterior)

### Supp. 9

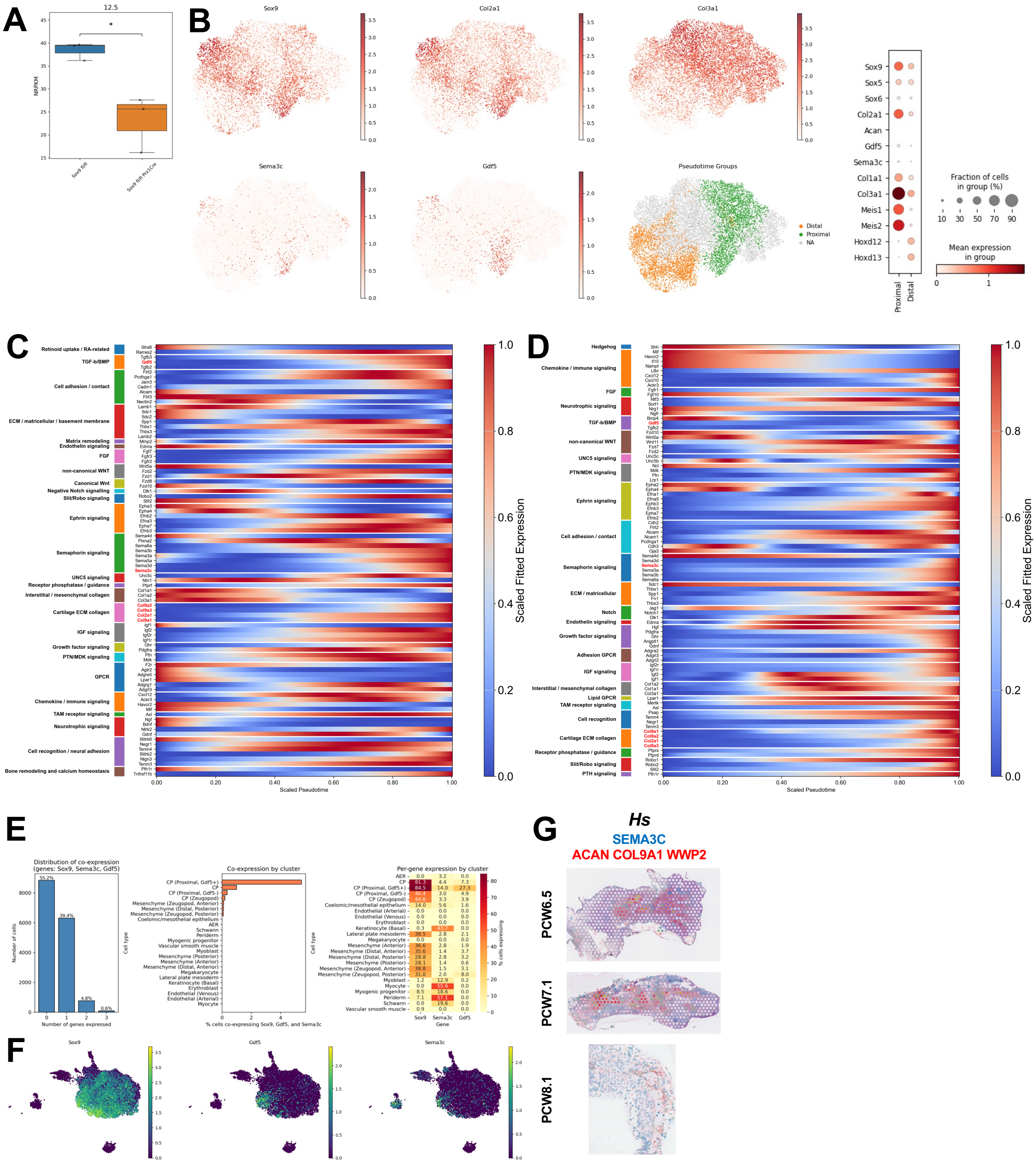

A
12.5
NRPKM
Sox9 fl/fl
Sox9 fl/fl Prx1Cre
B
Sox9
Col2a1
Col3a1
Sema3c
Gdf5
Pseudotime Groups
Distal
Proximal
NA
Sox9
Sox5
Sox6
Col2a1
Acan
Gdf5
Sema3c
Col1a1
Col3a1
Meis1
Meis2
Hoxd12
Hoxd13
Proximal
Distal
Fraction of cells in group (%)
Mean expression in group
C
Retinoid uptake / RA-related
TGF-b/BMP
Cell adhesion / contact
ECM / matricellular / basement membrane
Matrix remodeling
Endothelin signaling
FGF
non-canonical WNT
Canonical Wnt
Negative Notch signalling
Slit/Robo signaling
Ephrin signaling
Semaphorin signaling
UNC5 signaling
Receptor phosphatase / guidance
Interstitial / mesenchymal collagen
Cartilage ECM collagen
IGF signaling
Growth factor signaling
PTN/MDK signaling
GPCR
Chemokine / immune signaling
TAM receptor signaling
Neurotrophic signaling
Cell recognition / neural adhesion
Bone remodeling and calcium homeostasis
Scaled Pseudotime
Scaled Fitted Expression
D
Hedgehog
Chemokine / immune signaling
FGF
Neurotrophic signaling
TGF-b/BMP
non-canonical WNT
UNC5 signaling
PTN/MDK signaling
Ephrin signaling
Cell adhesion / contact
Semaphorin signaling
ECM / matricellular
Notch
Endothelin signaling
Growth factor signaling
Adhesion GPCR
IGF signaling
Interstitial / mesenchymal collagen
Lipid GPCR
TAM receptor signaling
Cell recognition
Cartilage ECM collagen
Receptor phosphatase / guidance
Slit/Robo signaling
PTH signaling
Scaled Pseudotime
Scaled Fitted Expression
E
Distribution of co-expression (genes: Sox9, Sema3c, Gdf5)
Number of cells
Number of genes expressed
Co-expression by cluster
% cells co-expressing Sox9, Gdf5, and Sema3c
Per-gene expression by cluster
Gene
% cells expressing
F
Sox9
Gdf5
Sema3c
G
Hs
SEMA3C
ACAN COL9A1 WWP2
PCW6.5
PCW7.1
PCW8.1

### Supp. 10

## A

| Dataset | Stage | Strain | Enrichment |
|---|---|---|---|
| Matsushita et al | E13.5 | Col2a1-cre | Col2a1-lineage enriched |
| Zhang et al | E12.5 | C57BL/6J | |
| Woods et al | E13.5, E15.5 | Prrx1-cre | |
| Song et al | E13.5-E14.5 | C57BL/6N | Soft tissues removed |
| Bian et al | E12.5-E15.5 | Gdf5-cre | Gdf5-lineage enriched |

## B

- CP
- CP (Proximal, Gdf5+)
- CP (Proximal, Gdf5-)
- CP (Zeugopod)
- Lateral plate mesoderm
- Mesenchyme (Anterior)
- Mesenchyme (Distal, Anterior)
- Mesenchyme (Distal, Posterior)
- Mesenchyme (Posterior)
- Mesenchyme (Zeugopod, Anterior)
- Mesenchyme (Zeugopod, Posterior)
- Chondrocyte
- Dermal fibroblast
- Early connective tissue
- IZ
- Mesenchyme (Pelvic girdle, Condensing)
- PHC
- Tendon/ligament progenitor (Distal Zeugopod)
- Tendon/ligament progenitor (Zeugopod)
- Fibroblast
- IZ (Chondrogenic)
- IZ (Talocrural)
- Mesenchyme
- Mesenchyme (Proximal)
- Perichondrium
- Pre-osteoblast
- Tendon/ligament progenitor
- Tendon/ligament progenitor (Patella)
- Tendon/ligament progenitor (Proximal)
- HC
- HC (Late)
- IZ (IA)
- Perichondrium (Inner)
- Perichondrium (Outer)
- Tendon/ligament progenitor (Patellar)

## C

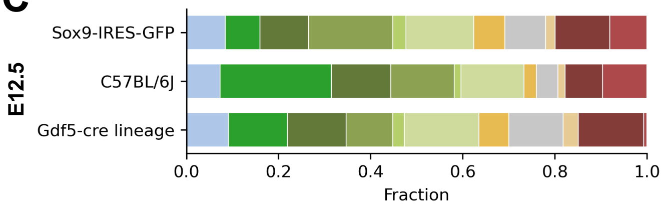


| | IZ |
|---|---|
| **Sox9-IRES-GFP** | 14.8% |
| **C57BL/6J** | 13.7% |
| **Gdf5-cre** | 16.2% |

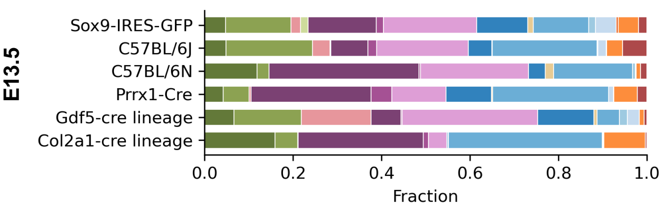


| | IZ | IZ (Chondrogenic) | IZ (Talocrural) |
|---|---|---|---|
| **Sox9-IRES-GFP** | 1.7% | 15.4% | 1.6% |
| **C57BL/6J** | 0.2% | 8.4% | 2% |
| **C57BL/6N** | 0% | 33.9% | 0.3% |
| **Prrx1-Cre** | <0.1% | 27.1% | 4.7% |
| **Gdf5-cre** | 0% | 6.8% | 0.3% |
| **Col2a1-cre** | <0.1% | 28.3% | 1.2% |

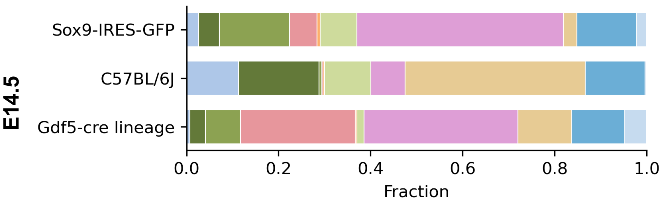


| | IZ |
|---|---|
| **Sox9-IRES-GFP** | 8% |
| **C57BL/6J** | 10.1% |
| **Gdf5-cre** | 1.5% |

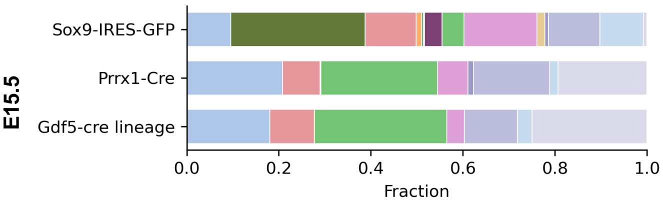


| | IZ | IZ (Chondrogenic) | IZ (IA) |
|---|---|---|---|
| **Sox9-IRES-GFP** | 0.2% | 3.8% | 4.8% |
| **Prrx1-Cre** | 0% | 0% | 25.3% |
| **Gdf5-cre lineage** | 0% | 0% | 28.7% |

#### Supp. 11

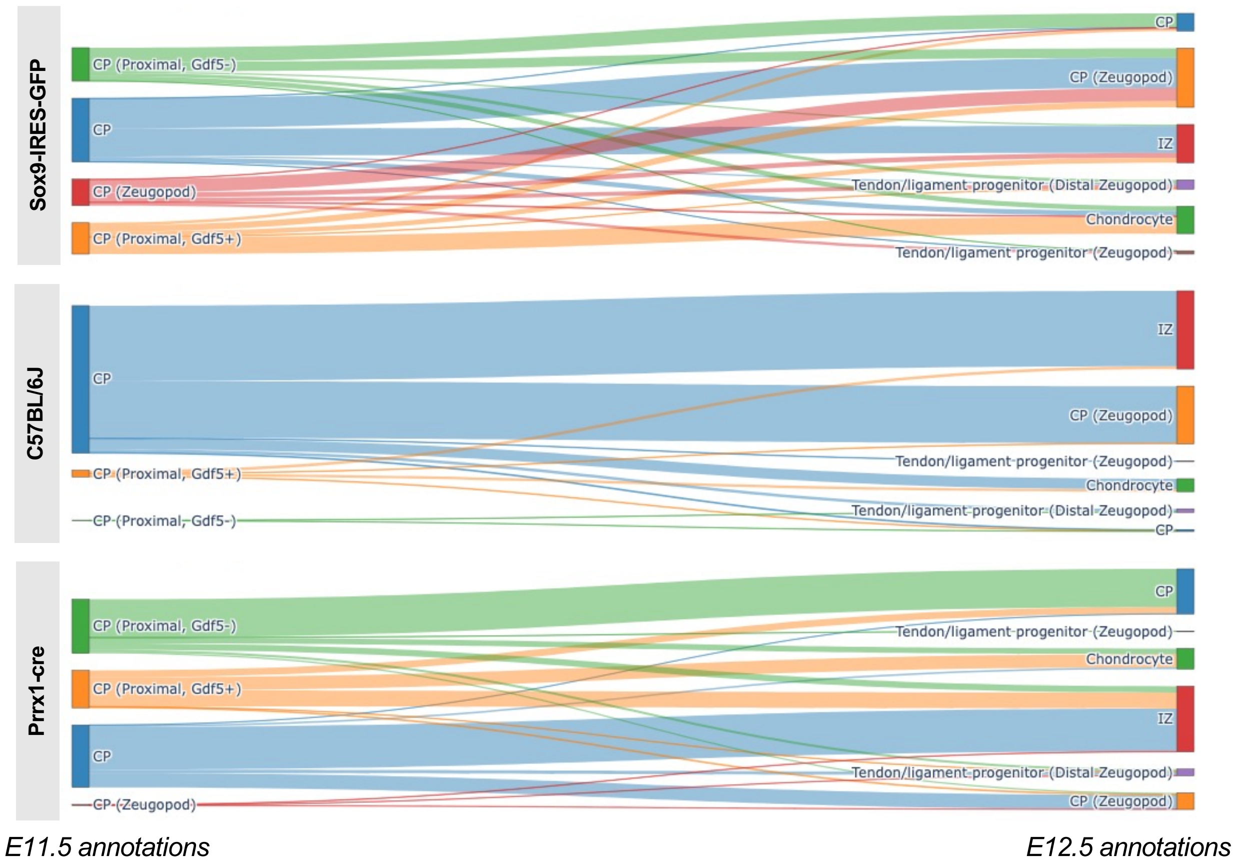

Sox9-IRES-GFP
CP (Proximal, Gdf5-)
CP
CP (Zeugopod)
CP (Proximal, Gdf5+)
CP
CP (Zeugopod)
IZ
Tendon/ligament progenitor (Distal Zeugopod)
Chondrocyte
Tendon/ligament progenitor (Zeugopod)
C57BL/6J
CP
CP (Proximal, Gdf5+)
CP (Proximal, Gdf5-)
IZ
CP (Zeugopod)
Tendon/ligament progenitor (Zeugopod)
Chondrocyte
Tendon/ligament progenitor (Distal Zeugopod)
CP
Prrx1-cre
CP (Proximal, Gdf5-)
CP (Proximal, Gdf5+)
CP
CP (Zeugopod)
CP
Tendon/ligament progenitor (Zeugopod)
Chondrocyte
IZ
Tendon/ligament progenitor (Distal Zeugopod)
CP (Zeugopod)
E11.5 annotations
E12.5 annotations

Supp. 12

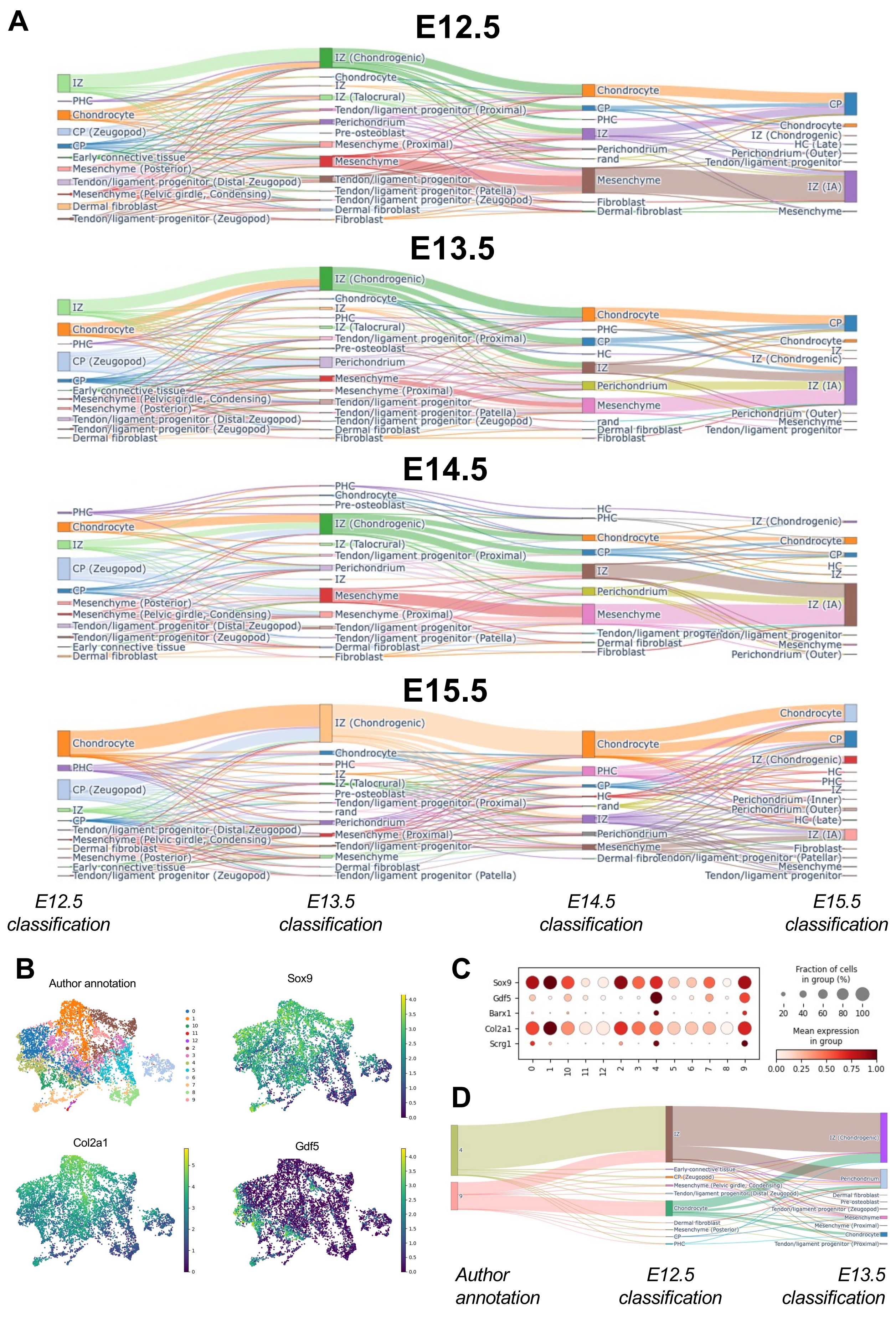
A
E12.5
E13.5
E14.5
E15.5
E12.5 classification
E13.5 classification
E14.5 classification
E15.5 classification
IZ
PHC
Chondrocyte
CP (Zeugopod)
CP
Early connective tissue
Mesenchyme (Posterior)
Tendon/ligament progenitor (Distal Zeugopod)
Mesenchyme (Pelvic girdle, Condensing)
Dermal fibroblast
Tendon/ligament progenitor (Zeugopod)
IZ (Chondrogenic)
IZ (Talocrural)
Tendon/ligament progenitor (Proximal)
Perichondrium
Pre-osteoblast
Mesenchyme (Proximal)
Mesenchyme
Tendon/ligament progenitor
Tendon/ligament progenitor (Patella)
Fibroblast
rand
HC
HC (Late)
Perichondrium (Outer)
Perichondrium (Inner)
IZ (IA)
Tendon/ligament progenitor (Patellar)
B
Author annotation
Sox9
Col2a1
Gdf5
C
Sox9
Gdf5
Barx1
Col2a1
Scrg1
Fraction of cells in group (%)
20 40 60 80 100
Mean expression in group
0.00 0.25 0.50 0.75 1.00
D
Author annotation
E12.5 classification
E13.5 classification

### Supp. 13

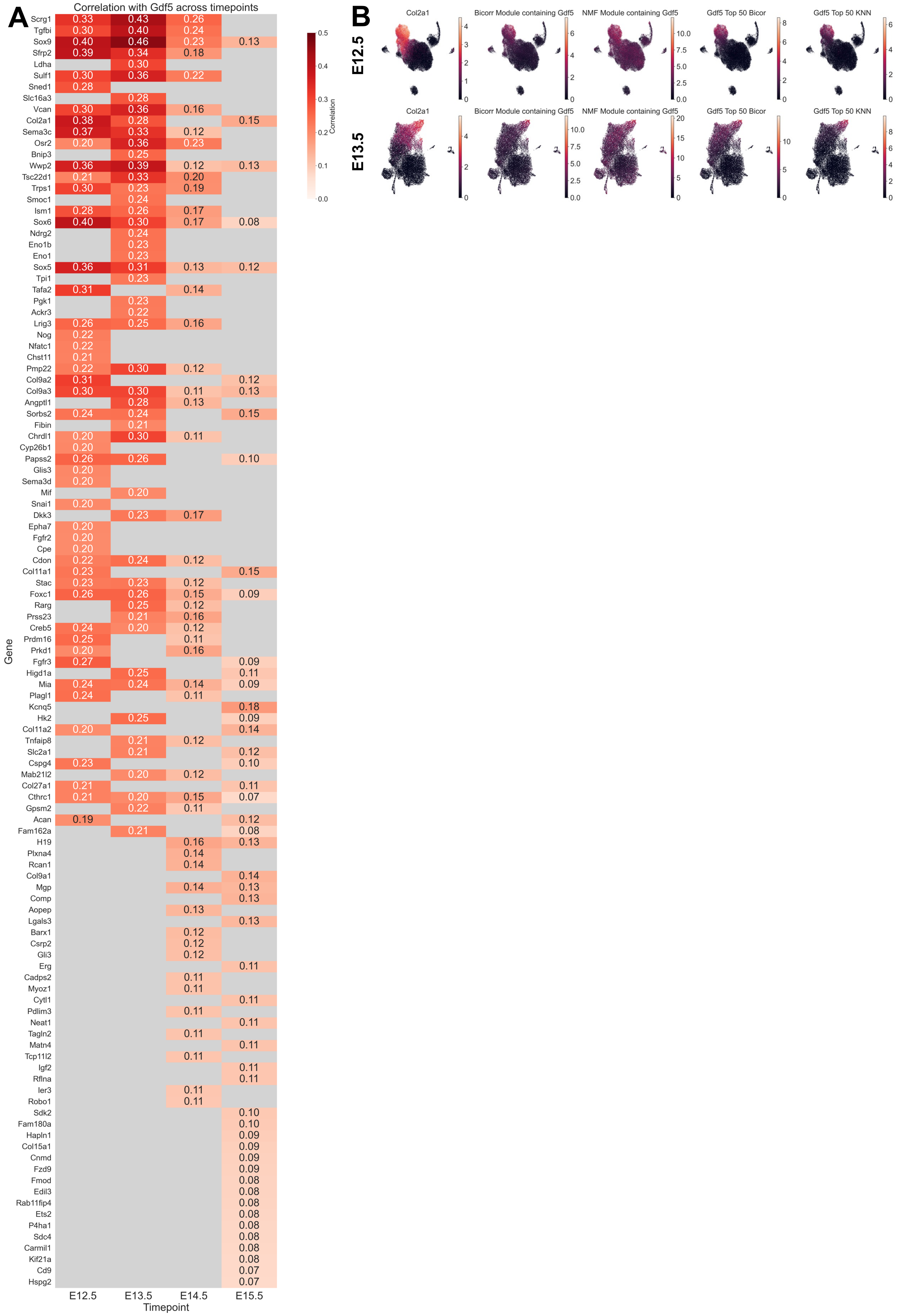
A
Correlation with Gdf5 across timepoints
Gene
Timepoint
E12.5
E13.5
E14.5
E15.5
Correlation
B
E12.5
E13.5
Col2a1
Bicorr Module containing Gdf5
NMF Module containing Gdf5
Gdf5 Top 50 Bicor
Gdf5 Top 50 KNN

**Supp. 14**

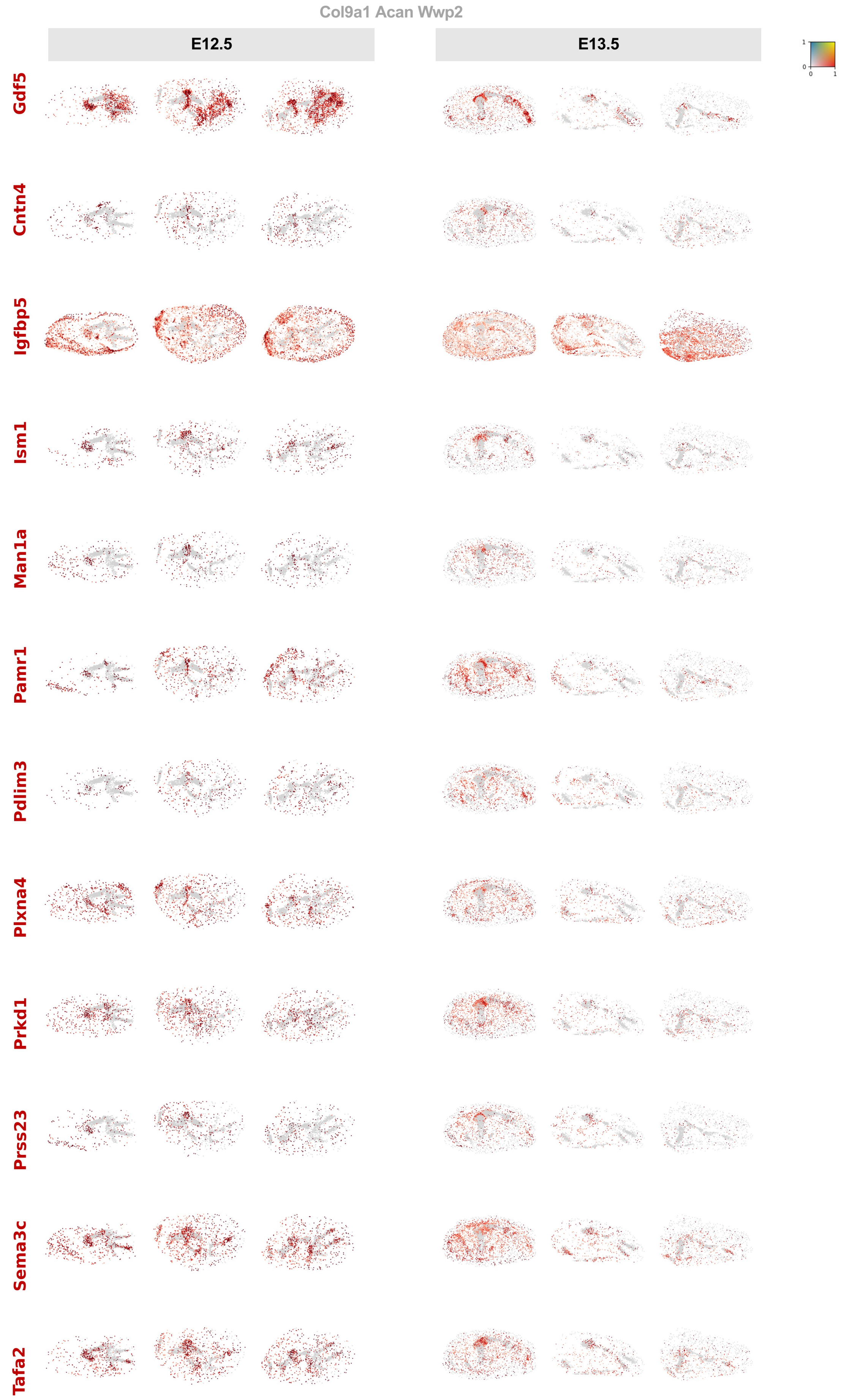
Col9a1 Acan Wwp2
E12.5
E13.5
1
0
0
1
Gdf5
Cntn4
Igfbp5
Ism1
Man1a
Pamr1
Pdlim3
Plxna4
Prkd1
Prss23
Sema3c
Tafa2

### Supp. 15

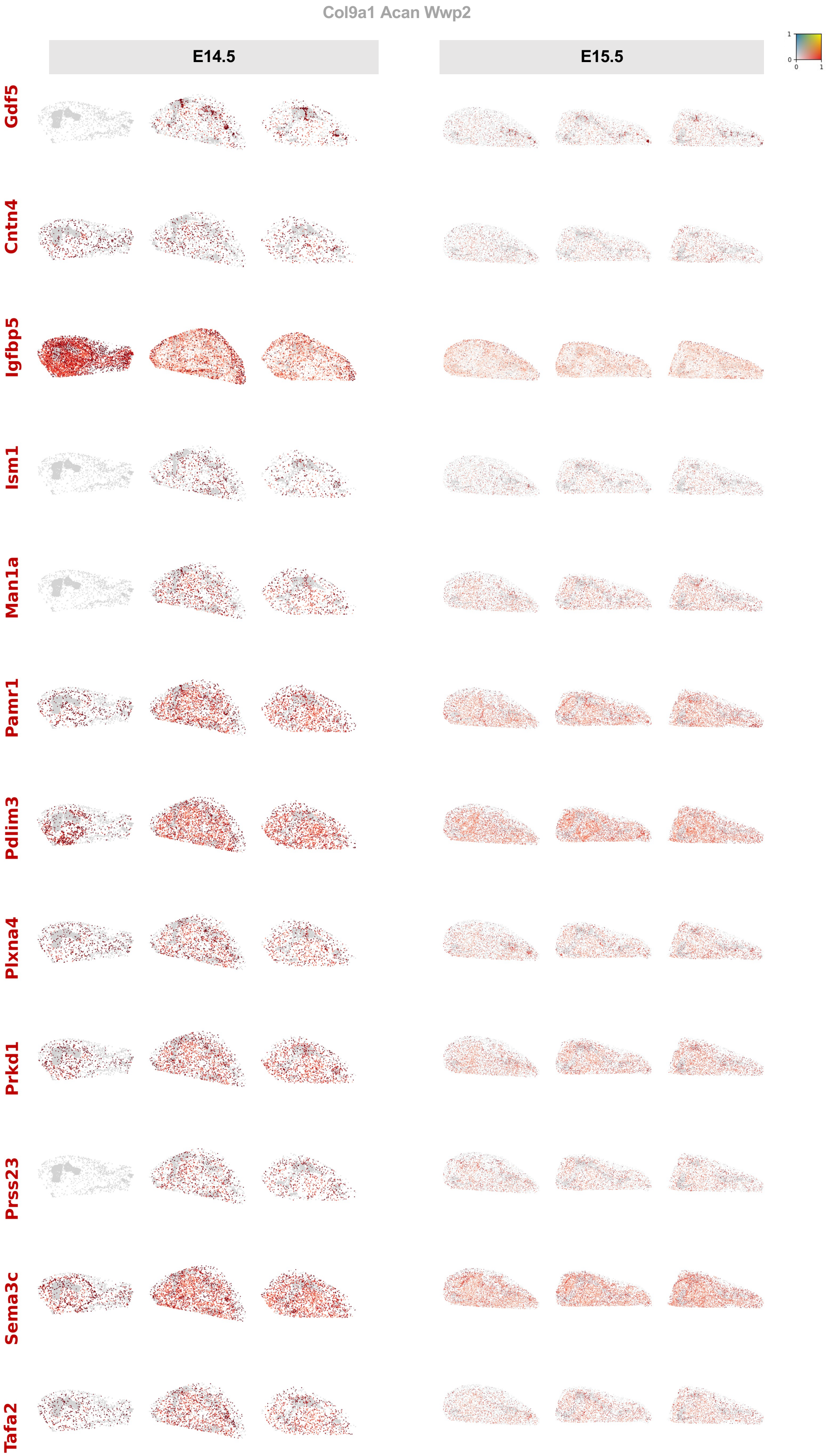

Col9a1 Acan Wwp2
E14.5
E15.5
1
0
0
1
Gdf5
Cntn4
Igfbp5
Ism1
Man1a
Pamr1
Pdlim3
Plxna4
Prkd1
Prss23
Sema3c
Tafa2

Supp. 16

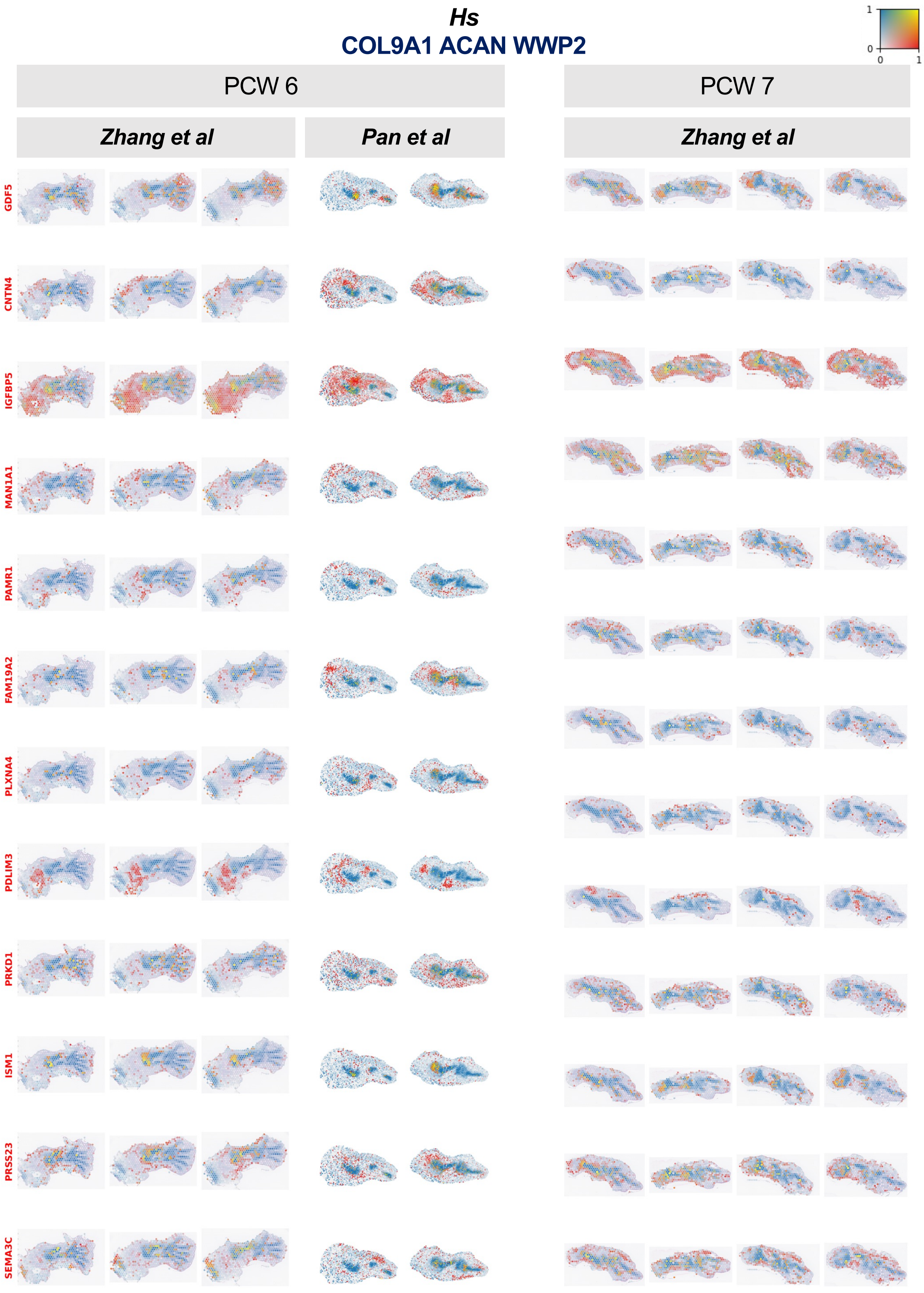
Hs
COL9A1 ACAN WWP2
1
0
0
1
PCW 6
PCW 7
Zhang et al
Pan et al
Zhang et al
GDF5
CNTN4
IGFBP5
MAN1A1
PAMR1
FAM19A2
PLXNA4
PDLIM3
PRKD1
ISM1
PRSS23
SEMA3C
